# Orthologs of an essential orphan gene vary in their capacities for function and subcellular localization in *Drosophila melanogaster*

**DOI:** 10.1101/2025.05.01.651694

**Authors:** Prajal H. Patel, Lars A. Eicholt, Andreas Lange, Kerry L. McDermott, Erich Bornberg-Bauer, Geoffrey D. Findlay

## Abstract

Orphan genes evolve rapidly, raising questions about whether their functions remain conserved or diverge across species. To address this, we investigated *goddard* (*gdrd*), an orphan gene essential for spermatogenesis in *Drosophila melanogaster*. Within the *Drosophila* genus, Gdrd proteins retain a conserved core structure but display substantial variation in length and primary sequence. Here we perform cross-species gene-swap assays in *D. melanogaster* testes to examine how these lineage-specific changes affect Gdrd function. Strikingly, the highly divergent *D. mojavensis* ortholog fully rescues fertility in *gdrd* null flies, suggesting that ancestral Gdrd acted within a conserved spermatogenesis pathway. By contrast, several orthologs, including one from a more closely related species, cannot substitute for the *melanogaster* gene. Cytological analysis shows that all divergent Gdrd orthologs retain some ability to interact with axonemes and ring centrioles, consistent with the protein’s structural conservation, but many non-complementing orthologs display weaker axonemal binding. Furthermore, all tested orthologs exhibit divergent localizations to organellar structures. Using computational analyses and molecular dynamics simulations, we identified intrinsic protein qualities that may account for several observations made in the gene swap assays. Rescuing orthologs bear motifs with shared physicochemical properties in their intrinsically disordered regions, while non-rescuing variants exhibit structural instabilities. Taken together, these findings show that while Gdrd’s ancestral structure and interactions are conserved, several orthologs have undergone lineage-specific evolutionary changes.

## INTRODUCTION

Orphan genes are phylogenetically restricted genes that encode proteins lacking detectable homology to known proteins (Tautz and Domazet-Lošo 2011; Zhao et al. 2024; Pereira et al. 2025). As a group, they comprise roughly 10–20% of eukaryotic genomes, with 30% as an upper limit (Khalturin et al. 2009; Prabh and Rödelsperger 2016). Because orphan genes lack well-studied homologs and characterized structural domains, their functions are difficult to predict (Middendorf et al. 2024). Nonetheless, many are implicated in species-specific adaptations involving behavior, reproduction, host-pathogen interactions, and speciation (Wilson et al. 2005; Khalturin et al. 2009; Fakhar et al. 2023). During their typically brief lifespans—encompassing gene birth, maturation, and eventual loss (Schmid and Aquadro 2001; Tautz and Domazet-Lošo 2011; Palmieri et al. 2014; Iyengar and Bornberg-Bauer 2023; Grandchamp et al. 2024; Lebherz et al. 2024)—low selective constraints facilitate the rapid evolution of their protein-coding sequences (Domazet-Loso and Tautz 2003; Wissler et al. 2013). These changes may refine orphan protein functions and enable their integration into cellular pathways (Domazet-Loso and Tautz 2003; Peng and Zhao 2024).

The *Drosophila melanogaster* gene *goddard* (*gdrd*) was identified as an orphan gene that plays an essential role in male fertility (Zhang et al. 2010; Gubala et al. 2017; Heames et al. 2020; Lange et al. 2021; Peng and Zhao 2024). Although other origination mechanisms cannot be excluded due to the gene’s advanced age (>40 million years), Gdrd’s testis-biased expression and the lack of structurally similar homologs suggest a possible *de novo* origin from non-coding sequences (Gubala et al. 2017; Lange et al. 2021; Peng and Zhao 2024). Functional analyses indicate that *gdrd* most likely acts during spermatid elongation, when Gdrd achieves peak expression and localization at the polymerizing axoneme and the insect ring centriole (Lange et al. 2021). Loss of *gdrd* results in stalled spermatid cysts at late spermatid elongation and the failure to produce mature sperm. At present, the precise role *gdrd* plays at the axoneme or the insect ring centriole remains unknown.

*gdrd* appears to be phylogenetically restricted to the *Drosophila* genus (Gubala et al. 2017; Heames et al. 2020; Lange et al. 2021; Peng and Zhao 2024). Within the *melanogaster* species group (Fig. 1a), it has evolved under purifying selection, indicating functional conservation (Gubala et al. 2017; Peng and Zhao 2024). In more distantly related *Drosophila* species, however, *gdrd* orthologs only weakly align with the *D. melanogaster* protein using BLASTP (Gubala et al. 2017; Lange et al. 2021). Despite these divergences in primary amino acid sequence, all Gdrd orthologs maintain their structural composition: a central helix of conserved length and flanking N- and C-terminal intrinsically disordered regions (IDRs) (Fig. 1a) (Lange et al. 2021). The main structural variation among extant orthologs is gene length, as orthologs in distant species often contain substantially longer IDRs at their termini (Lange et al. 2021). Whether these differences in gene length or primary sequence affect protein function remains an open question.

**Fig. 1.**
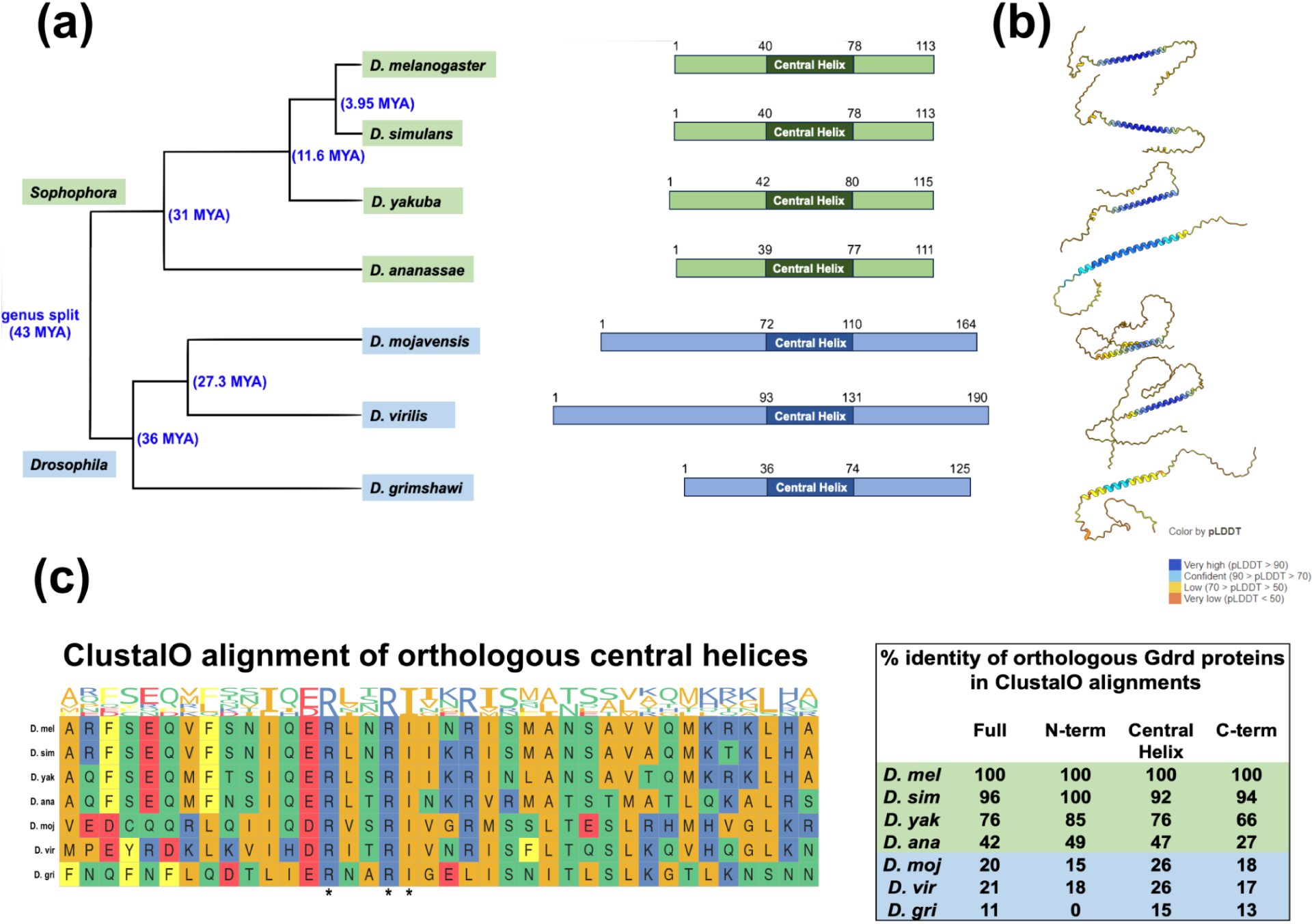
Gdrd orthologs from the *Drosophila* subgenus exhibit amino acid sequence divergence and coding sequence lengthening at both the N- and C- termini. a) *Left: gdrd* orthologs are present throughout the *Drosophila* genus within both the *Drosophila* and *Sophophora* subgenera, which diverged around 43 million years ago (MYA). Divergence times (in blue brackets and font) were estimated using TimeTree. *Middle:* The Gdrd orthologs share a structure and can be divided into regions of disordered N-and C-termini and a central helix. *Sophophora* subgenus *gdrd* genes (green) typically have shorter coding sequences compared to their *Drosophila* subgenus (blue) counterparts. b) The central helix is conserved in all Gdrd orthologs and predicted with very high confidence by AlphaFold2 (>90 pLDDT). See bottom right for pLDDT score legend. c) *Left:* ClustalO multiple sequence alignment of the central helix region. Three conserved amino acid residues common to all seven orthologs are marked with asterisks. *Right:* Analysis of amino acid conservation within ClustalO multiple sequence alignment using percent identity as a metric. The table shows percent identity conservation between *D. melanogaster* Gdrd and orthologs over the proteins’ full lengths and by protein region. As expected, distantly related orthologs exhibit decreased amino acid conservation, with the *D. grimshawi* ortholog presenting the greatest sequence divergence. The disordered *D. grimshawi* Gdrd N-terminus shows no detectable primary sequence homology to the *D. melanogaster* protein.

Interestingly, multiple studies have identified IDRs as a defining feature of *de novo*-emerged orphan genes (Ángyán et al. 2012; Landry et al. 2015; Basile et al. 2017; Heames et al. 2020; Eicholt et al. 2022; Heames et al. 2023; Aubel et al. 2024; Middendorf and Eicholt 2024; Peng and Zhao 2024). Although the precise functions of these regions remain largely unresolved, IDRs generally provide essential protein characteristics such as structural flexibility and accessible interaction sites (Brown et al. 2011). Because IDR properties are primarily physicochemical (e.g., charge, polarity) and largely independent of amino acid position, these regions can tolerate frequent substitutions, insertions, and deletions with minimal impact on function (Uversky 2017; LeBlanc et al. 2024; Jemth 2025). While this primary sequence mutability may mimic evolution under relaxed selection, IDR physicochemical properties remain subject to evolutionary constraint (Brown et al. 2011). Consequently, functionally important physicochemical motifs within IDRs persist across time (Zarin et al. 2021) and can be identified using position-independent, alignment-free homology tools (Chow et al. 2024).

As orphan genes evolve, rapid sequence divergence can have varied consequences, ranging from negligible effects on protein–protein interactions (Vakirlis et al. 2020; Chen et al. 2024; Middendorf et al. 2024) to the refinement of existing functions or even the emergence or loss of function. Because the functions and interaction partners of many orphan genes remain poorly characterized, their evolutionary trajectories are difficult to resolve (Bornberg-Bauer and Eicholt 2026). Here, we perform assessments of *gdrd* ortholog function and localization in *D. melanogaster* in combination with AlphaFold2 (AF2) predictions, molecular dynamics (MD) simulations, physicochemical analyses of IDRs, and sequence homology analyses to address two questions. Do amino acid changes within a conserved structure alter protein function, or are such substitutions largely inconsequential? And, what kinds of functional refinements has the gene undergone during its evolutionary history?

Consistent with Gdrd’s structural conservation (Fig. 1a) across the genus, we find that a highly diverged ortholog can functionally replace the *D. melanogaster* gene, suggesting that the ancestral Gdrd protein was integrated into a spermatogenesis-related pathway. In several descendant lineages, however, amino acid changes have led to overall structural destabilization and altered physicochemical properties within terminal IDRs. These changes correlate with an inability to function in *D. melanogaster*. Taken together with observations that several orthologs exhibit distinct subcellular localization patterns when expressed in *D. melanogaster* testes, these findings suggest that Gdrd has retained, lost, and acquired different physicochemical and localization properties during its evolution across the genus.

## RESULTS

### Gdrd orthologs from the *Drosophila* subgenus show substantial primary sequence divergence from the *melanogaster* protein

Using TBLASTN, we searched sequenced Drosophilid genomes for orthologs of *gdrd* and a syntenically associated gene, *dnaaf6*. This analysis corroborates the previously published evolutionary history of *gdrd* (Gubala et al. 2017), demonstrating that the gene is syntenic with *dnaaf6* throughout the *Drosophila* genus—including both the *Drosophila* and *Sophophora* subgenera—while being absent from the *willistoni* sublineage of *Sophophora*. In total, we identified 33 and 66 *Drosophila* and *Sophophora* subgenus orthologs, respectively.

Multiple sequence alignments of all identified orthologs using Clustal Omega (ClustalO) (Sievers et al. 2011) reveal that the central helices of these orthologs are largely alignable, with the notable exception of *D. gunungcola* (Figure S1). The N- and C-terminal regions of the orthologous proteins, however, show lineage-specific alterations in length (Fig. 1a). As the size of the ancestral gene at the base of the genus is presently unknowable, it is uncertain if these termini have expanded or contracted in extant *gdrd* genes. Orthologs present outside of the *melanogaster* species group (subgenus *Sophophora*) have slightly longer C-termini (Figures 1a, S1). Indeed, increases in *gdrd* gene sizes in the *obscura* species group (subgenus *Sophophora*) appear to be dependent on increased C-termini (Figure S1). By contrast, the gene size increases observed in many sublineages of the *Drosophila* subgenus, which can reach up to twice the size of *melanogaster* group orthologs, are due principally to the presence of longer N-termini (Figure S1). Interestingly, the largest *gdrd* orthologs are present in the *virilis* and *melanica* species groups (Figure S1). These orthologs encode proteins that contain regions enriched with glycine and aspartic acid residues, which are not present in related *repleta* species group orthologs (Figure S1). Altogether, these data indicate that the lengths of Gdrd N- and C-termini vary in a lineage-specific manner across the *Drosophila* genus.

To investigate the evolutionary fates of *gdrd* orthologs present within the *Drosophila* genus, we selected orthologs from the *Sophophora* and *Drosophila* subgenera (Fig. 1a), which split ∼40-43 million years ago (MYA) (Russo et al. 1995; Kumar et al. 2017). This included four related orthologs from the subgenus *Sophophora* (inclusive of the *melanogaster* gene) and three divergent orthologs from the subgenus *Drosophila* (Fig. 1a). AF2 predictions confirm the structural conservation of all seven orthologous proteins (Fig. 1b).

To evaluate amino acid sequence conservation among divergent *gdrd* orthologs, we performed pairwise BLASTP comparisons between the *D. melanogaster* protein and the selected *Drosophila* genus orthologs. As expected, *Sophophora* subgenus orthologs are alignable across their full lengths, with sequence identities ranging from 46–96% (Figure S2). These levels of sequence identity indicate reliable homology <u>(Rost 1999)</u> and are also observed in pairwise comparisons across a larger set of *melanogaster* species group orthologs (Figure S3). By contrast, the three orthologs from the *Drosophila* subgenus exhibit limited sequence homology (0-38%) over only partial alignments (Figure S2). When analyzing BLASTP alignments by protein region, the *D. melanogaster* protein’s similarity with *D. mojavensis* and *D. virilis* Gdrd is confined to the central helix, which appears to be the most conserved region among all Gdrd orthologs (Figure S2, S3). Notably, the *D. grimshawi* ortholog exhibits no detectable similarity with the *D. melanogaster* protein when the central helix is analyzed separately (Figure S2). Pairwise BLASTP analyses between *Drosophila* subgenus species orthologs and the *melanogaster gdrd* termini detects no significant homology within any of the selected *Drosophila* subgenus species. This homology detection failure likely results from highly divergent sequences and lengths of termini in these orthologs, combined with the intrinsic disorder characteristic of these regions (Figure S2, S3).

To get around these sequence homology detection issues, we performed a multiple sequence alignment of the selected *gdrd* orthologs using ClustalO. Among *Sophophora* subgenus orthologs, full-length pairwise alignments with *D. melanogaster* Gdrd shows 42–96% identity (Fig. 1c). This range decreases to 15-21% identity for *Drosophila* subgenus orthologs, with *D. grimshawi* being the least conserved. When percent identity was examined by protein region (N-terminus, central helix, and C-terminus), we found broadly similar levels of amino acid conservation across Gdrd orthologs, with occasional cases (*D. ananassae* and *D. grimashawi*) in which the terminal regions exhibited greater sequence divergence than the central helix (Fig. 1c).

Collectively, these data indicate that Gdrd orthologs across the *Drosophila* genus have retained their overall structure but also experienced lineage-specific amino acid sequence changes. This divergence may stem from lineage-specific adaptations arising through coevolution with other proteins, from reduced selective constraints on the gene, or both.

### Expression of codon-optimized *gdrd* orthologs in *D. melanogaster*

Cross-species complementation assays offer a powerful framework for testing the functional consequences of protein divergence and have been used previously in *Drosophila* evolution studies (Bayes and Malik 2009; Saint-Leandre et al. 2020; Brand and Levine 2022). To examine how sequence divergence influences *gdrd* function, we performed gene swap experiments using a previously described *gdrd* rescue construct that both restores fertility to *gdrd* null (full gene deletion) mutants and expresses C-terminally hemagglutinin (HA)-tagged proteins amenable to subcellular localization studies using immunohistochemistry (Lange et al. 2021).

To promote consistent expression in *D. melanogaster*, we codon-optimized each *gdrd* ortholog and integrated all constructs into the same genomic locus via phiC31-mediated recombination before crossing them into a *gdrd* null background. Codon optimization, which alters synonymous sites, has been successfully applied to *Drosophila* orthologs in previous studies (Saint-Leandre et al. 2020; Brand and Levine 2022). Because synonymous substitutions can influence gene expression (Zhang and Qian 2025), optimization could theoretically affect transcript abundance. To test this, we assessed transcript levels using RT-PCR and confirmed robust and comparable transcript abundance across nearly all constructs used in this study (Figure S4).

We initiated our swap analyses by replacing the wild-type *D. melanogaster gdrd* coding sequence in the rescue construct with a codon-optimized version and assessing its expression in testes. To facilitate developmental staging and imaging, we assessed Gdrd expression and subcellular localization in isolated cysts. Each cyst consists of descendants from a single germline stem cell division and two somatically derived cyst cells that enclose the developing germ cells. Within each cyst, germ cells progress synchronously through spermatogenesis, beginning with coordinated mitotic divisions during spermatogonial phases, pre-meiotic growth during spermatocyte stages, meiosis, and then several post-meiotic spermiogenic stages, which include spermatid elongation initiation, early and late elongating spermatids, spermatid individualization, and spermiation of mature sperm (see graphical overview of spermatogenesis in Figure S5) (Fabian and Brill 2012). Because Gdrd localizes to the basal body, transition zone, and axoneme during spermatid elongation (Lange et al. 2021), we examined its expression relative to Unc, an early marker of centriole-to-basal body conversion (Baker et al. 2004). We find that *D. melanogaster* Gdrd colocalizes with transition zone marker Unc and that the codon-optimized construct faithfully reproduces prior expression and localization patterns observed with the original *gdrd* rescue (Figure S5). Detailed analysis of Unc and Gdrd localization patterns during spermatogenesis are presented in the Supplementary Text.

### The Gdrd central helix is sufficient for axoneme and transition zone localization

Sequence analyses revealed that the disordered N- and C-termini of Gdrd orthologs vary in length (*D. mojavensis*, *D. virilis*) and amino acid composition (*D. ananassae*, *D. grimshawi*) (Fig. 1a,c). To assess the functional relevance of these terminal IDRs and contextualize their evolutionary divergence, we performed structure-function analyses using modified *D. melanogaster* rescue constructs lacking either terminal region. The N-terminal truncation (GdrdΔN) removes 36 amino acids after the initial methionine, and the C-terminal truncation (GdrdΔC) removes the final 35 amino acids (Fig. 2a). MD simulations based on AF2 modelings predict that both truncated proteins remain structurally stable and retain their central α-helices (Fig. 2b). Each construct was then introduced into *D. melanogaster* and assessed for functional complementation. The full-length codon-optimized construct restores fertility to wild-type levels, indistinguishable from the non–codon-optimized rescue (n = 25 per genotype, unequal variance *t*-test, *P* = 0.08) (Fig. 2c). By contrast, both truncations fail to rescue fertility, indicating that the N- and C-terminal regions are essential for Gdrd function (n = 25 per genotype, unequal variance *t*-test, *P* < 0.0001) (Fig. 2c).

**Fig. 2.**
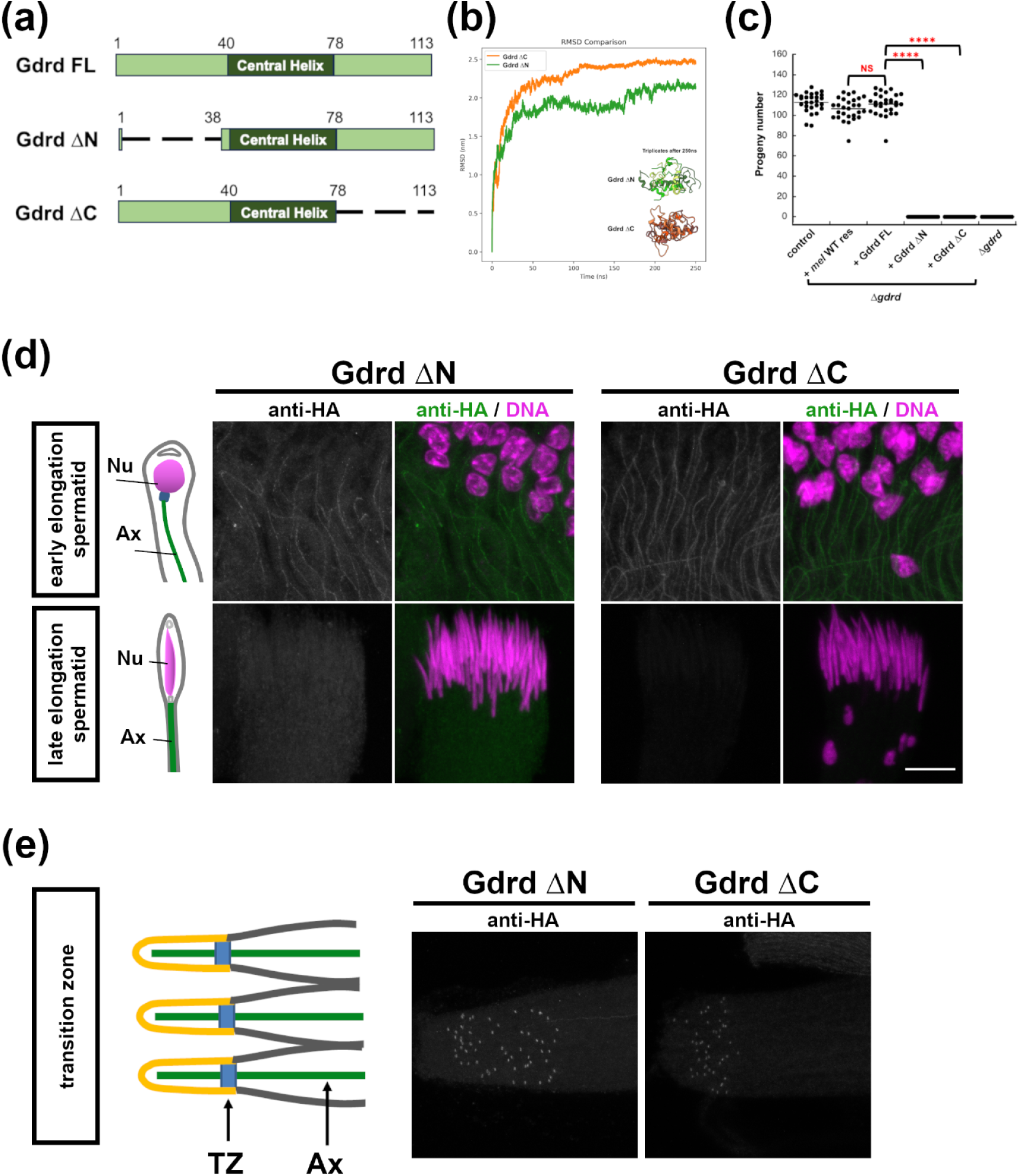
Gdrd’s central helix mediates localization to axonemes and transition zones. **a)** Diagrams of full length codon-optimized *D. mel* Gdrd (Gdrd FL), *D. mel* Gdrd N-terminal truncation (Gdrd ΔN), and *D. mel* Gdrd C-terminal truncation (Gdrd ΔC). b) Averaged backbone RMSD values from triplicate 250 ns MD simulations of Gdrd ΔN and Gdrd ΔC indicate stable folding of the central helix. Data for each run are available in supplementary materials (Fig. S6). c) Neither Gdrd truncation can complement the *gdrd* null fertility defect. Statistical test: *t*-tests with unequal variance. NS = not significant, **** = *P* < 0.0001. d) □N and □C Gdrd proteins localize to axonemes in both early and late elongation spermatid cysts. Diagrams depict nuclear shaping criteria used in developmental staging early and late elongating spermatids (Nu = nucleus; Ax = axoneme). e) Both □N and □C Gdrd proteins localize to transition zones. The diagram illustrates distal ends of mature cysts (TZ, transition zone; Ax, axoneme). Ciliary cap (yellow) compartmentalizes the polymerizing end of the axoneme. Scale bar = 10 μm in all images.

To determine whether the inability of the N- and C-terminally truncated Gdrd proteins to rescue fertility could be caused by altered protein expression, we performed western blot analysis on whole testes. Despite robust transcript levels (Figure S4), both truncated proteins are difficult to detect on western blots, suggesting either instability or reduced solubility due to the truncation of the terminal IDRs (Figure S7). Indeed, previous studies have demonstrated the role IDRs play in protein solubility (Tretyachenko et al. 2017; Uversky 2017; Borcherds et al. 2021; Eicholt et al. 2022). We next examined the expression and localization patterns of truncated Gdrd protein using immunohistochemistry. While we can detect both proteins using immunofluorescence, their signals are weak when compared to the full length protein (Fig. 2d, Figure S5). Furthermore, the C-terminally truncated protein appears the least stable and becomes undetectable in late elongating spermatid cysts (Fig. 2d). The diminished levels of both truncated proteins may account for the inability of either construct to restore fertility to *gdrd* mutant males. Nevertheless, even at low protein levels, both truncated proteins localize to axonemes in early and late elongating spermatids (Fig. 2d) and are detectable at the transition zone (Fig. 2e). Altogether, these observations suggest that Gdrd’s central helix, which is common to both constructs, mediates its localization to the axoneme and insect ring centriole.

### A *gdrd* ortholog from the *Drosophila* subgenus is functional in *D. melanogaster*, while other orthologs of lesser or equal divergence are not

We next replaced the *D. melanogaster gdrd* coding sequence in the rescue construct with codon-optimized orthologs from *D. simulans*, *D. yakuba*, *D. ananassae*, *D. mojavensis*, *D. virilis*, and *D. grimshawi*. To determine if these orthologous proteins can be exogenously expressed in *D. melanogaster* testes, we performed western blot analysis on testis protein extracts and immunohistochemistry on whole-mount testes. Because the orthologs differ in size, amino acid composition, and net charge, they likely differ in their electrophoretic mobilities during western blot transfer, making it difficult to compare protein levels directly. The *D. simulans*, *D. yakuba*, *D. mojavensis*, and *D. virilis* orthologs, however, are readily detectable on blots, while the *D. ananassae* and *D. grimshawi* proteins are only weakly detectable (Fig. 3A), suggestive of diminished stability *in vivo* and/or weaker electrophoretic mobility during blotting.

**Fig. 3.**
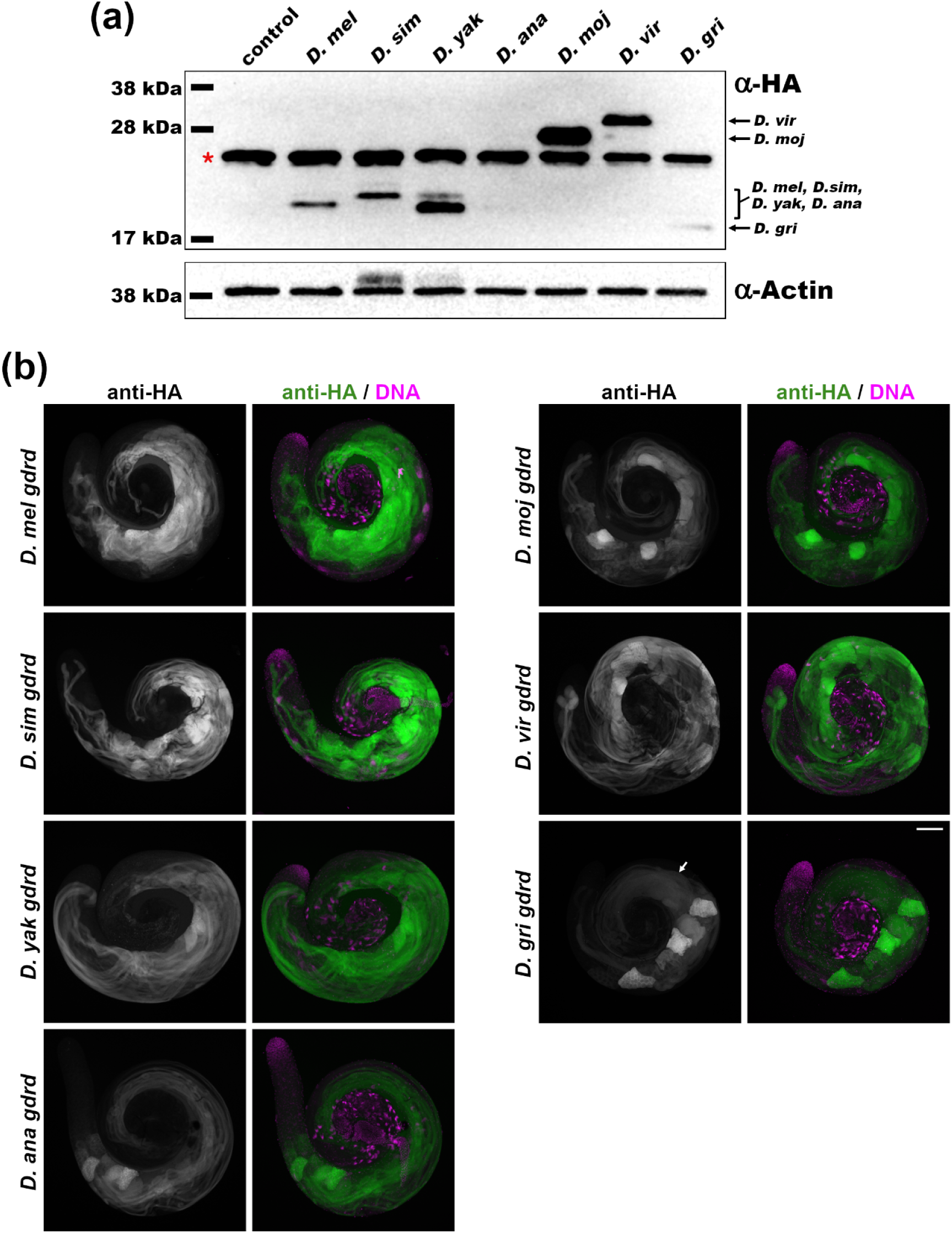
Exogenous expression of *gdrd* orthologs in *D. melanogaster* testes appears largely robust. a) Western blot of lysates from *D. melanogaster* testes expressing HA-tagged *gdrd* orthologs. With the exception of *D. ananassae* (*D. ana*) and *D. grimshawi* (*D. gri*) Gdrd proteins, most orthologs are stably expressed. Control is the *gdrd* mutant genetic background. Red asterisk labels a non-specific band present in both control and experimental lysates. Actin serves as loading control. b) Spatial and temporal expression of Gdrd orthologs in *D. melanogaster* testes reflect wild-type *melanogaster* expression patterns. *D. melanogaster* testes were labeled with anti-HA (gray scale or green). The apical ends of the testes, which contain the youngest cysts, are on the top left side of each image, while the basal ends containing the nuclei of older cysts are at the center. The anti-HA signal intensity of *D. melanogaster* (*D. mel*), *D. simulans* (*D. sim*), and *D. yakuba* (*D. yak*) orthologs appears robust. This signal appears diminished in more phylogenetically distant orthologs: *D. ananassae* (*D. ana*), *D. mojavensis* (*D. moj*), *D. virilis* (*D. vir*), and *D. grimshawi* (*D. gri*). Within late elongating *D. grimshawi* spermatid cysts (arrow), the anti-HA signal appears diffuse, suggesting that the *D. grimshawi* protein doesn’t localize to axonemes. Nuclei are labeled with DAPI (magenta). Scale bar = 100 μm.

Using immunohistochemistry on whole testes, we found that all exogenously introduced *gdrd* orthologs are detectable (Fig. 3B). As the ortholog gene swap constructs are driven by the *D. melanogaster* gene’s regulatory elements, inclusive of the 5’ and 3’ UTRs, their spatiotemporal expression patterns in testes closely parallel that of the native gene, with maximal protein accumulation in post-meiotic round spermatids (Fig. 3B). However, relative to the *D. melanogaster* protein, immunohistochemical signals for the *D. ananassae*, *D. mojavensis*, *D. virilis*, and *D. grimshawi* orthologs appear weaker, suggesting decreased stability, altered localization, or both (Fig. 3). Notably, the *D. grimshawi* ortholog displayed a faint and diffuse signal in both early and late elongating spermatid cysts, indicating that the *D. grimshawi* protein also fails to associate with axonemes (Fig. 3, Figure S8a,b). Altogether, these data indicate that Gdrd orthologs can be expressed in *D. melanogaster* testes albeit with variable stabilities or detectabilities.

To evaluate the functional conservation of *gdrd* orthologs, we tested their ability to rescue fertility defects in *gdrd* mutants. Successful complementation was interpreted as evidence of functional conservation. Negative results in these assays, however, can have several non-mutually exclusive explanations, including loss or diminished functional capabilities, functional divergence, or lineage-specific co-evolution with binding partners.

Codon-optimized *D. melanogaster gdrd* restores fertility to wild-type levels (*n* = 30 per genotype, unequal variance *t-*test, *P* = 0.25) (Fig. 4a). Although *Sophophora* Gdrd orthologs share sequence and structural conservation (Fig. 1a-c), they exhibit distinct capacities to restore fertility in *gdrd* mutants. Unsurprisingly, the *simulans gdrd* ortholog, which shares 96% identity with the *D. melanogaster* ortholog, fully restores fertility (*n* = 30, unequal variance *t-*test, *P* = 0.55) (Fig. 4a). Despite sharing 76% amino acid identity with *D. melanogaster*, the *D. yakuba* ortholog restores only 4–43% of wild-type fertility levels across replicates (*n* = 30 per genotype for each replicate, unequal variance *t*-tests, *P* < 0.0001) (Fig. 4a, Figure S9). The more divergent *D. ananassae* ortholog (42% sequence identity) consistently fails to restore any fertility to *gdrd* mutants (*n* = 30, unequal variance *t-*test, *P* < 0.0001) (Fig. 4a). Potentially, this reduced rescue capacity may reflect suboptimal expression of this ortholog in *D. melanogaster* testes. Altogether, these data suggest that even when both sequence and structure are fairly well conserved, two *Sophophora* subgenus *gdrd* orthologs cannot fully compensate for loss of the *D. melanogaster* gene.

**Fig. 4.**
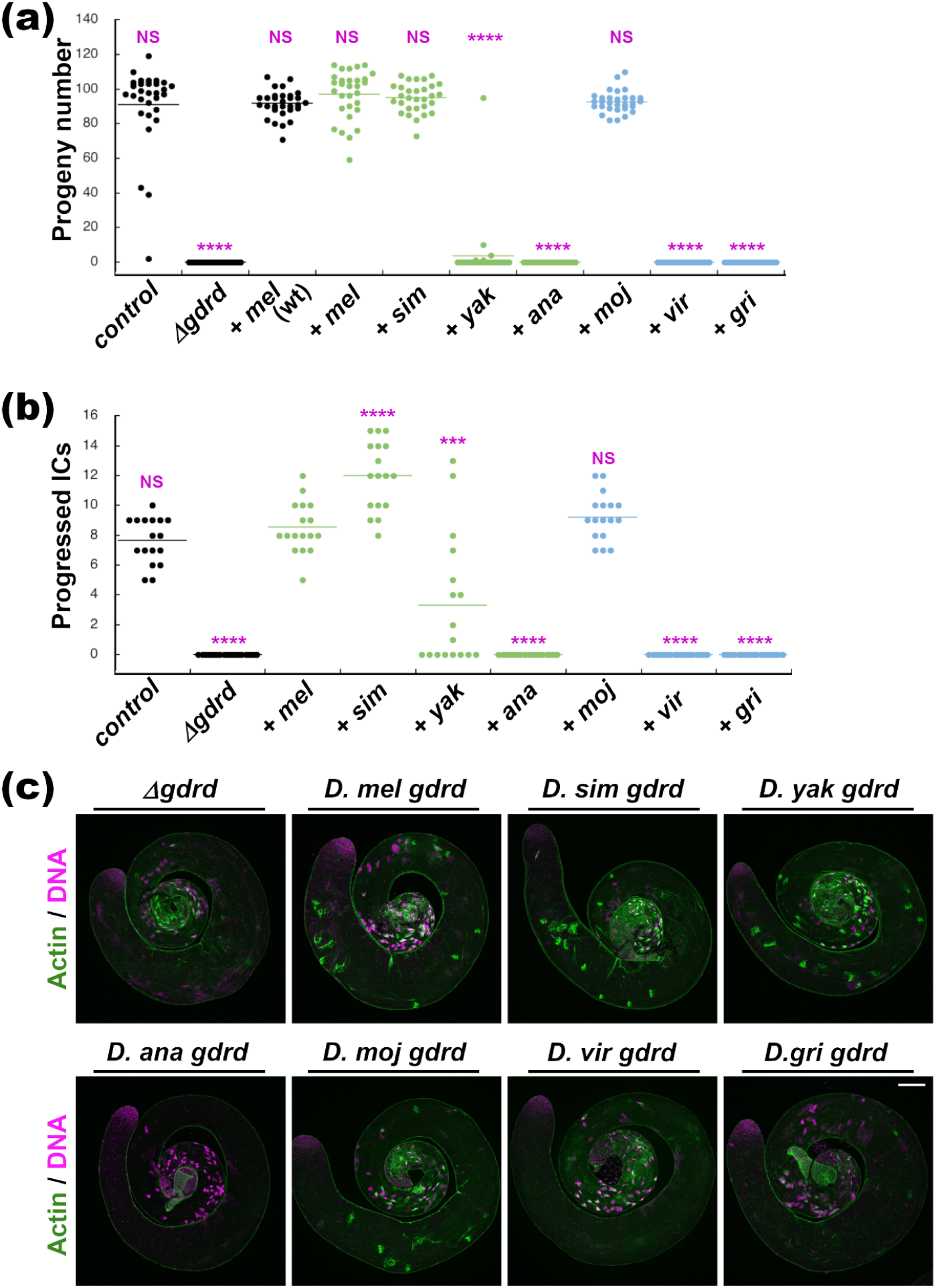
A *gdrd* ortholog that diverged over 40 million years ago fully restores fertility to *gdrd* null *D. melanogaster* males, while several other orthologs of equal or lesser divergence do not. a) Fertility assay to assess the function of *gdrd* orthologs in *D. melanogaster*. *Δgdrd* mutants expressing codon-optimized *D. melanogaster* (*mel*) Gdrd exhibit similar fertility levels when compared with either control flies (*w^11118^*) *or Δgdrd* mutants expressing the original, non-codon optimized *D. melanogaster* rescue (*mel wt*) (*P* = 0.25 and 0.08, respectively). *Δgdrd* mutants expressing codon-optimized *D. melanogaster* (*mel*) also exhibit similar fertility levels when compared with *Δgdrd* mutants expressing the *D. simulans* (*sim*) or *D. mojavensis* (*moj*) orthologs (*P* = 0.55 and 0.14 respectively). The *D. yakuba* ortholog (*yak*) appears only partially functional (mean progeny number = 3.7; *P* < 0.0001). The expression of *D. ananassae* (*ana*), *D. virilis* (*vir*), or *D. grimshawi* (*gri*) orthologs fails to rescue *Δgdrd* male sterility (*P* < 0.0001 in all cases). Statistical test: *t*-tests with unequal variance. NS = not significant, **** = *P* < 0.0001. b) Testes from either non-rescuing (*ana*, *vir, gri*) or poorly rescuing (*yak*) ortholog gene swaps display either no or diminished progressed ICs, respectively, suggesting a defect in initiating spermatid individualization. The partial restoration of IC progression in *yak*-expressing testes is consistent with the partial rescue observed in the fertility assay (panel a). Two-sample *t*-tests with unequal variances show significantly reduced levels of IC progression for the *ana*, *vir*, and *gri* orthologs (all *P* < 0.0001) and the *yak* ortholog (*P* = 0.00015), and significantly increased numbers of progressed ICs for the *sim* ortholog (*P* < 0.0001). In the graph, NS = not significant, *** = *P* < 0.001, **** = *P* < 0.0001. Non-parametric statistical analysis of the data in panels (a) and (b) gave equivalent results (see Supplemental Text). c) In *gdrd* mutants, ICs form but do not translocate along the elongated spermatid cysts. Progressing ICs are readily observable in fertile or partially fertile ortholog gene swaps as determined by phalloidin labeling of actin-rich ICs (green). *D. ananassae* gene swap testes, however, often lack IC formation altogether, indicating that exogenous expression of the *D. ana* ortholog enhances the *gdrd* mutant phenotype. DAPI labels DNA (magenta). Scale bar = 100 μm.

Strikingly, the highly divergent *gdrd* orthologs present within the *Drosophila* subgenus also exhibit variable abilities to restore fertility in *D. melanogaster gdrd* mutants. Much to our surprise, the *D. mojavensis* ortholog (20% amino acid sequence identity) fully restores fertility to wild-type levels (n = 30, unequal variance *t-*test, *P* = 0.14) (Fig. 4a). This finding has several implications. First, it suggests that the protein is functional in *D. melanogaster* despite considerable amino acid sequence divergence (Fig. 1c). Second, because the common ancestor of *D. mojavensis* and *D. melanogaster* lies at the base of the *Drosophila* genus, the ancestral *gdrd* gene likely already performed the core functions found in extant *D. melanogaster*. Third, the lengthening of the terminal IDRs in *D. mojavensis* (Fig. 1a) appears to have little impact on the protein’s central functions (Fig. 4a). On the other hand, the similarly divergent orthologs from *D. virilis* (21% identity) and *D. grimshawi* (11% identity) (Fig. 1c) fail to restore fertility (n = 30, unequal variance t-tests, *P* < 0.0001) (Fig 4a). While reduced protein expression or stability could be a factor in the inability of the *D. grimshawi* ortholog to restore fertility, the *D. virilis* ortholog is expressed at similar levels to the *D. mojavensis* protein and, at least as judged by western blot, at higher levels than the *D. melanogaster* ortholog (Fig. 3). Thus, within the context of these gene swap assays in *D. melanogaster*, amino acid sequence divergence has diminished the functional capabilities of some *gdrd* orthologs while being inconsequential in others.

Because *gdrd* loss-of-function mutations disrupt sperm individualization (Lange et al. 2021), we examined this process in gene-swapped testes by quantifying progressed individualization complexes (ICs). Gene swap testes expressing non-complementing orthologs had no progressed ICs, indicating complete loss of function in this assay (Fig. 4b). In contrast, *gdrd* mutant testes expressing the *D. yakuba* ortholog, which partially restores fertility, exhibit progressing ICs, though in significantly reduced numbers compared with those expressing the *D. melanogaster* protein (*D. melanogaster*: 8.5 ± 0.4; *D. yakuba*: 3.3 ± 1.1; *n* = 15, unequal variance *t*-test, *P* < 0.0001) (Fig. 4b). Interestingly, *D. simulans* ortholog gene swap testes show an increased number of progressing ICs despite restoring fertility to wild-type levels (Fig. 4b). By contrast, *gdrd* mutant testes expressing the *D. ananassae* ortholog exhibit more severe defects than *gdrd* null mutants. Whereas IC formation is reduced but detectable in *gdrd* mutants (Lange et al. 2021), most testes expressing the *D. ananassae* ortholog fail to form ICs altogether, suggesting that spermatid elongation never reaches completion (Fig. 4c). Because this phenotype appears in two independently generated insertion strains, it is unlikely to result from secondary mutations. Thus, expression of the *D. ananassae* ortholog in *D. melanogaster* may interfere with spermatogenic processes, thereby attenuating elongation. To test whether this effect is dominant or dominant negative, we measured fertility in flies carrying one copy each of the *D. ananassae* and *D. melanogaster* orthologs in a *gdrd* mutant background. These flies displayed fertility levels comparable to controls (Figure S10), indicating that the *D. ananassae* ortholog acts recessively and that the attenuation effect is mild.

### Orthologous Gdrd proteins tend to retain axonemal and insect ring centriole localization but also exhibit divergent subcellular localization patterns

A potential explanation for diminished function in non-rescuing *gdrd* orthologs might be disrupted protein localization. To examine this possibility, we observed the associations of HA-tagged orthologous Gdrd proteins with either the axoneme or the transition zone/ insect ring centriole. As the *D. grimshawi* protein shows weak expression (Fig. 3) and neither axonemal or transition zone localizations (Figure S8), we exclude this ortholog in the remaining analyses.

As previously shown (Figure S5), codon-optimized *D. melanogaster* Gdrd decorates the axoneme in early elongating spermatids, a stage marked by extensive axoneme elongation but limited nuclear re-modeling. This localization persists through late elongating spermatid stages and disappears in individualizing spermatid cysts (Figure S11b,d). The *D. simulans, D. yakuba* and *D. mojavensis* Gdrd orthologs, which restore fertility either fully or partially, show robust axonemal localization (Fig. 5b). By contrast, the *D. virilis* and *D. ananassae* orthologs, which fail to rescue fertility, localize weakly to axonemes (Fig. 5b). Overall, we find that all orthologs retain some axonemal localization, but those with diminished axonemal binding correspondingly fail to restore fertility in *gdrd* mutant flies.

**Fig. 5.**
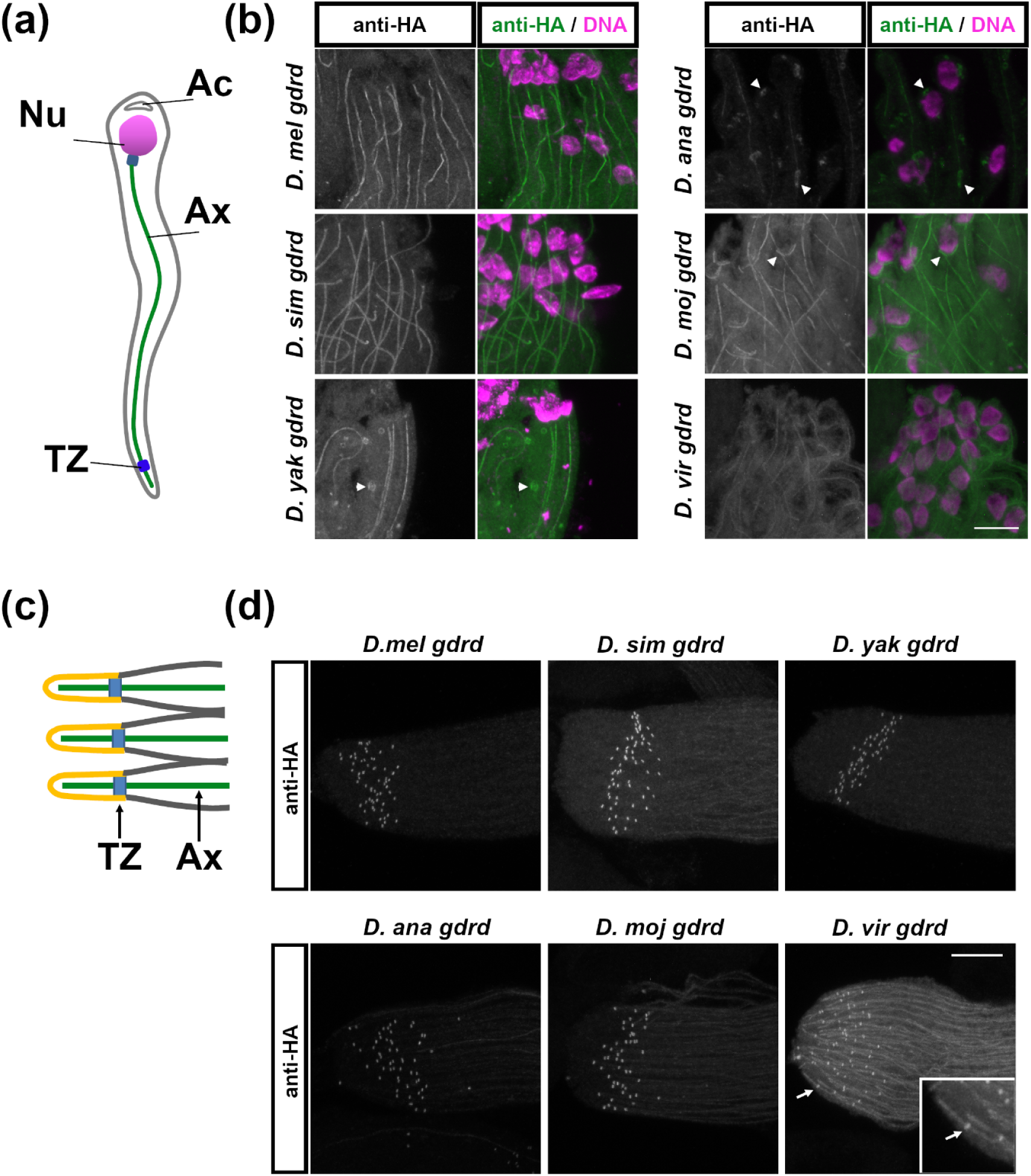
Gdrd orthologs show variable localization to axonemes and often exhibit divergent subcellular localization patterns in elongating spermatids. **a)** Diagram of early elongation spermatid (Nu, nucleus; Ax, axoneme; Ac, acrosome; TZ, transition zone). b) In early elongation spermatids, HA-tagged Gdrd orthologs localize to axonemes in all rescuing and partially rescuing gene swaps (*D. mel*, *D. sim*, *D. yak*, and *D. moj*). In non-rescuing *D. vir* and *D. ana* ortholog gene swaps, however, axonemal localization is reduced or greatly diminished. Some orthologs also exhibit divergent localizations at this stage (arrowheads). The *D. yak* ortholog decorates a round organellar structure. The *D. ana* ortholog associates with structures that abut the spermatid nuclei or axonemes, and the *D. moj* protein localizes to the apical surface of the round nuclei (arrowhead). The basal and apical ends of cysts are at the tops and bottoms of the images, respectively. c) Diagram depicts the distal end of a spermatid cyst (Ax, axoneme; TZ, transition zone). d) All Gdrd orthologs localize to the transition zone. The *D. vir* ortholog breaches the ciliary gate (inset). Scale bar = 10 μm.

We next examined if orthologous Gdrds also localize to the transition zone. As the distal end of spermatid cysts cannot be developmentally staged, we cannot make inferences about localization strength. Interestingly, all tested orthologs maintain localization to the transition zones, including the *D. ananassae* protein, which shows the most diminished localization at axonemes (Fig. 5d). Notably, the *D. virilis* protein associates with the axoneme within the ciliary cap (Fig. 5d). This suggests that the *D. virilis* Gdrd might have evolved a novel subcellular localization pattern. Alternatively, the presence of *D. virilis* Gdrd within the ciliary cap could reflect a loss of ciliary gate function inherent to the *D. melanogaster gdrd* mutant.

Several tested orthologs also display subcellular localization patterns that differ markedly from those of the *D. melanogaster* protein. These differences are not restricted to elongating spermatids, where Gdrd is thought to function, but are often apparent earlier in spermatogenesis, including in primary spermatocytes and in spermatids initiating elongation. In both *D. virilis* and *D. mojavensis* gene swaps, the corresponding Gdrd orthologs localize to the nuclear periphery in spermatocytes (Figure S12b) and early spermatids (Fig. 5b; Figure S13b). Notably, *D. mojavensis* Gdrd is enriched along the apical region of the nuclear envelope, resembling proteins associated with the tubulin-associated nuclear cap (Fig. 5b; Figure S13b).

In contrast, the *D. ananassae* Gdrd ortholog localizes to structures reminiscent of the Golgi apparatus in spermatocytes (Figure S12b) and subsequently to the Golgi-derived acroblast during elongation initiation (Figure S13b) and early spermatid elongation (Figure 5b). Intriguingly, both *D. simulans* and *D. yakuba* Gdrd proteins localize near the acrosome at the basal end of spermatid nuclei (Figure S11b), with the *D. yakuba* Gdrd additionally associating with a round organellar structure in late round spermatids (Figure 5b). Besides these nuclear and golgi/acroblast/acrosome localizations, orthologs also exhibit additional localization patterns. The *D. simulans* and *D. grimshawi* Gdrds transiently associate with the mitochondrial derivative and spermatid plasma membrane, respectively, during elongation initiation (Figure S13b). Taken together, these observations raise the possibility that divergent Gdrd orthologs might have evolved interactions with additional partners or affinities for different subcellular compartments.

To the extent that orthologous protein functions and localizations can be assessed in *D. melanogaster* cells, these data collectively suggest that, despite structural conservation, amino acid sequence divergence among Gdrd orthologs has major effects on both these properties.

### Gdrd orthologs exhibit sublineage-specific alterations in physicochemical composition of terminal IDRs

Amino acid sequence divergence can potentially lead to changes in a protein’s intrinsic properties. Because our AF2 predictions, structure-function analyses, and localization studies highlight the structural and functional conservation of the central helix, we next examined whether amino acid sequence divergence within the terminal IDRs contributes to diminished function or altered localization in ortholog gene swaps. As highly disordered regions evolve rapidly due to weak structural constraints, we applied k-mer–based, alignment-free tools, SHARK-Dive (Chow et al. 2024) and SHARK-Capture (Chow et al. 2025), to identify conserved motifs based on physicochemical similarity. These analyses show that the *D. mojavensis* termini more closely resemble those of *Sophophora* orthologs (Fig. 6a). The *D. mojavensis* N terminus shares a motif with *Sophophora* subgenus orthologs (PLDTSED in *Sophophora*; DTTHRSED and PISESIETG in *D. mojavensis*; Fig. 6a), while this motif is absent in *D. grimshawi* and *D. virilis* orthologs, which do not functionally rescue in *D. melanogaster* (Fig. 6b). Additionally, the *D. mojavensis* ortholog’s C-terminus has a substantially higher normalized SHARK-Dive score when compared to *D. melanogaster* than all other *Drosophila* subgenus orthologs analyzed, including closely related *repleta* group species (Fig. S14).

**Fig. 6.**
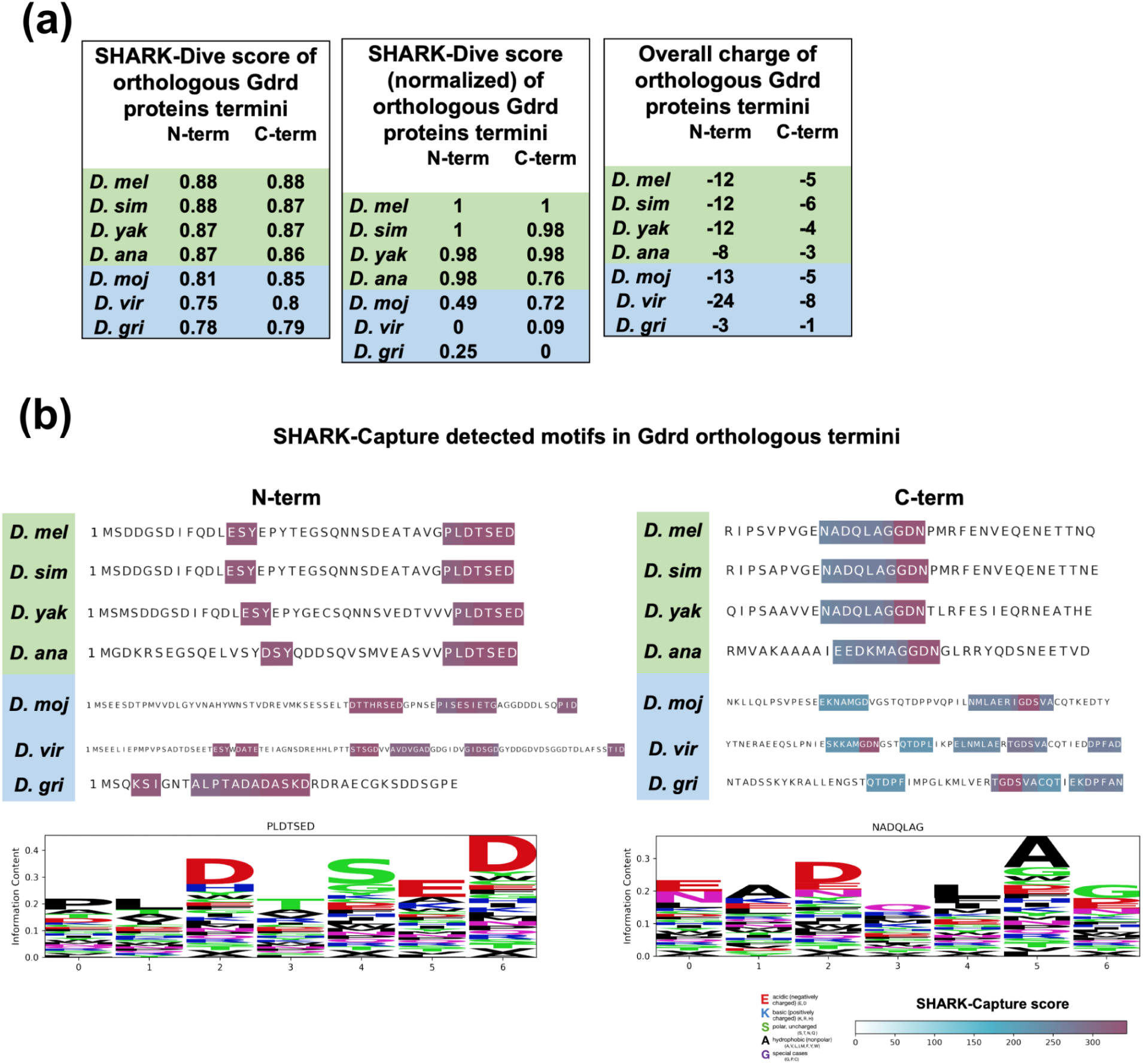
Alignment-free and charge analysis reveal that *D. mojavensis* terminal IDRs resemble those of *Sophophora* group species. a) Raw (*left*) and normalized (*middle*) SHARK-Dive scores and overall ionic charge (*right*) for the N- and C-terminal regions of all tested orthologs. For both metrics, the *D. mojavensis* termini are more similar to their respective termini of *Sophophora* subgenus orthologs than to more closely related subgenus *Drosophila* orthologs. *D. virilis* and *D. grimshawi* termini differ considerably from those of *Sophophora*. b) Motifs identified by SHARK-Capture show that the *D. mojavensis* Gdrd N- and C-termini harbor terminal motifs that are similar in physicochemical pattern to those found in Sophophora ortholog termini (top). Sequence logos summarize the most frequent consensus motif class detected across ortholog termini (bottom), illustrating conservation via property-preserving substitutions rather than strict residue identity. The bottom-right legend indicates SHARK similarity scores and the amino-acid property scheme used to assess motif similarity.

In addition to the identification of conserved physicochemical motifs, we also examined variation in net charge within the N- and C-terminal regions of Gdrd orthologs across the *Drosophila* genus. Gdrd orthologs within several *Drosophila* subgenus lineages have N-termini with higher negative charges compared to orthologs present in *Sophophora* (Figure S15). Within the *virilis* and *melanica* species groups, this is most likely due to the presence of aspartic acid/glycine rich regions in these orthologs (Figure S1b). Interestingly, the net charges of the *D. mojavensis* termini mirror that of *Sophophora* orthologs, whereas the *D. grimshawi* and *D. virilis* termini differ strongly (Fig. 6a, Figure S15). Collectively, these similarities between the *D. mojavensis* and *D. melanogaster* orthologs may contribute to the ability of the *D. mojavensis* ortholog to restore fertility in *gdrd* mutant males.

### MD simulations show that Goddard’s conserved central α-helix remains stable across lineages, while the N- and C-termini remain disordered

Since the non-rescuing *D. virilis* ortholog and the rescuing *D. mojavensis* ortholog differ primarily in the strength of their localizations to the axoneme and insect ring centriole (Fig. 5b,d), it is possible that divergence of the terminal IDRs (Fig. 6) may modulate Gdrd’s binding affinity and/or stability at these subcellular sites. To determine whether an ortholog’s ability to function in *D. melanogaster* correlates with predicted structural stability, we evaluated the stability of AlphaFold2-predicted protein structures (Fig. 1b) through molecular dynamics (MD) simulations conducted in triplicate over 250 ns. Previous studies have shown that structural predictions for orphan proteins can yield implausible results (Aubel et al. 2023; Liu et al. 2023; Middendorf and Eicholt 2024), and MD simulations can test the stability of the predicted structures (Lange et al. 2021; Middendorf et al. 2024; Peng and Zhao 2024).

Our modeling and MD simulations revealed that all orthologous Gdrd proteins exhibit structural flexibility (Fig. 7). Notably, we consistently observe flexible terminal regions characterized by disorder and transient short helix formations (Fig. 7a). In contrast, the central helical region remained largely stable across all simulations (Figure 7). Root Mean Square Deviation (RMSD) analyses confirm this intrinsic flexibility through high deviation values indicative of dynamic behavior (Seffernick et al. 2019) (Fig. 7b, left). Among the studied orthologs, *D. ananassae* had the highest RMSD values after relaxation, suggesting it is the most flexible or least stable, followed by *D. grimshawi* and *D. virilis*, while *D. mojavensis* appeared the most stable (Fig. 7c, top left). Interestingly, western blot analysis of exogenously expressed *D. ananassae* and *D. grimshawi* in *D. melanogaster* testes supports decreased stability (Fig. 3a). On the other hand, the *D. virilis* ortholog, along with related orthologs within the *virilis* and *melanica* species, bear an aspartic acid/glycine rich region (Figure S1) that may confer increased flexibility.

**Figure 7:**
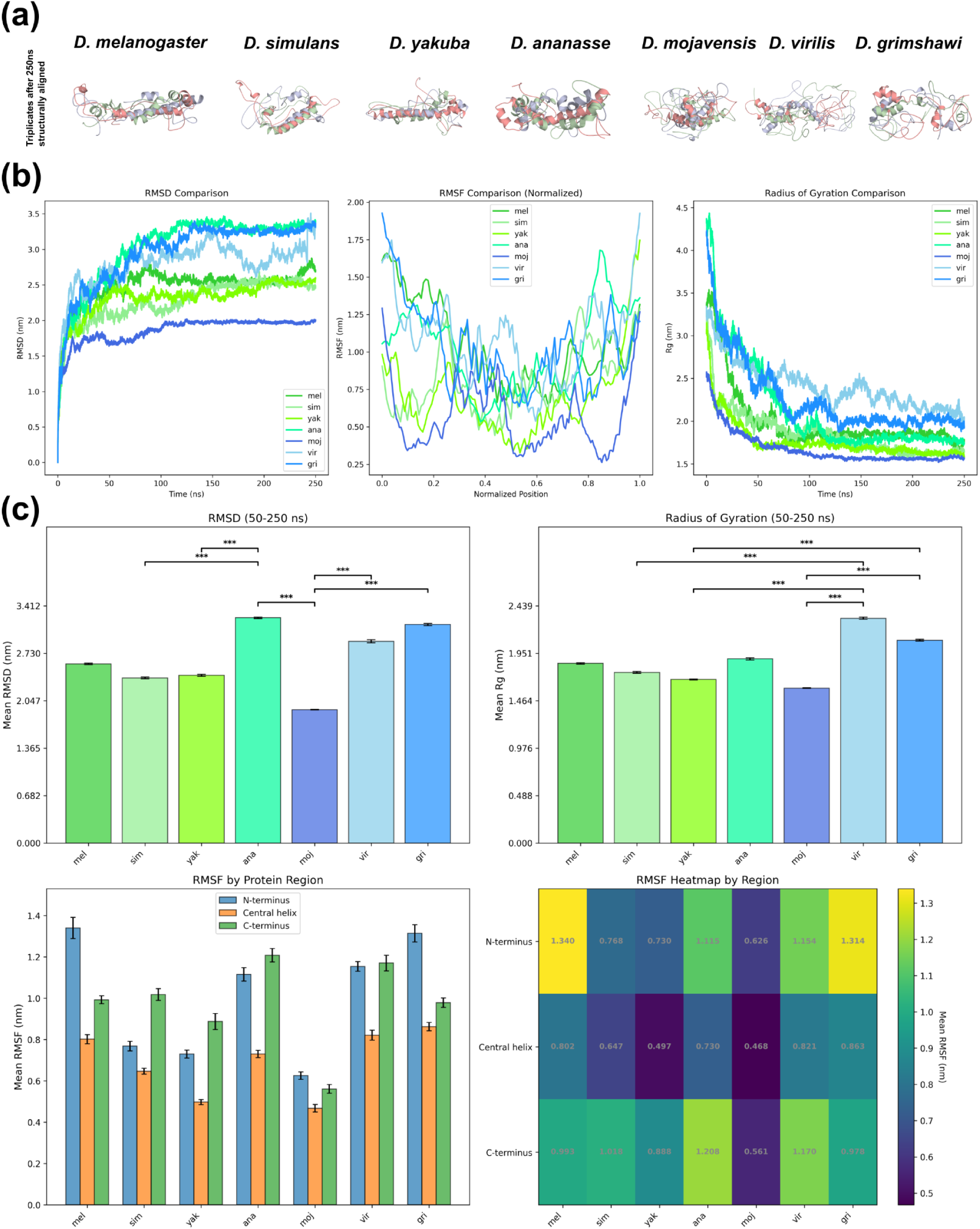
MD simulations of Gdrd orthologs over 250ns. a) Structural alignments of triplicate MD simulations after 250 ns highlight the dynamic structures of Gdrd orthologs, while the central helix remains stable. b) Averaged backbone RMSD over time (*left*), α-C RMSF per normalized residue position (*center*) and radius of gyration over time (*right*) of all Gdrd orthologs. *D. ananassae* shows by far the highest RMSD, indicating relative instability. α-C RMSF is high at the termini for all orthologs, in accordance with their disorder, while central residues fluctuate less and remain a structured helix overall. The trend of radii of gyration shows that all orthologs become more compact over time, likely forming molten globule-like structures. c) Mean RMSD, radius of gyration, and RMSF values averaged over the equilibrated 50–250 ns region. Kruskal–Wallis H-tests detected significant interspecies differences (*** = p < 0.001), followed by Dunn’s post-hoc tests with Bonferroni correction for 21 pairwise comparisons (see Supplement). Only the top 5 most significant comparisons are shown here. *D. ananassae* exhibited the highest RMSD, whereas *D. virilis* showed the largest radius of gyration. Flexibility was highest in the termini, while the conserved central helix remained stably folded across all orthologs. Single runs and α-C RMSF per residue available in supplementary materials (Figure S16).

An examination of residue-specific flexibility using Root Mean Square Fluctuation (RMSF) analysis (Fig. 7b, center) confirms that the stable central helix exhibits substantially fewer fluctuations than the disordered termini. The N-terminus of the *D. melanogaster* ortholog exhibits the highest fluctuations during simulations, while the *D. mojavensis* protein shows the least fluctuations along the whole protein (Fig. 7c, bottom). Additionally, we used Radius of Gyration (Rg) analysis to determine the overall compactness and stability across orthologs (Fig. 7b, right). Consistent with RMSD and RMSF observations, orthologs display decreasing radii of gyration over simulation time (Fig. 7b), indicative of initial structural relaxation followed by stable yet flexible conformations, characteristic of molten globule-like structures. After initial relaxation (first 50 ns), the *D. virilis* and *D. grimshawi* proteins display a significantly higher mean Rg than other orthologs (Fig. 7c, top right). Collectively, our results show that Gdrd orthologs retain a conserved, stably folded core but differ in the flexibility and compaction of their disordered termini, suggesting the hypothesis that functional divergence could have arisen through evolutionary fine-tuning of terminal IDRs rather than wholesale structural changes.

## DISCUSSION

### Ancestral Gdrd at the base of the genus was most likely functional

A central question regarding *gdrd* is when the gene acquired functionality, given that identifiable orthologs are currently restricted to the *Drosophila* genus (Fig. 1, Figure S1). Our current findings suggest that the gene was most likely functional at the base of the genus. Several pieces of evidence support this conclusion. First, even with significant sequence changes that hinder homology based detection, Gdrd shows structural conservation across the genus (Fig. 1a), indicating the existence of structural constraints required for the maintenance of function. Second, we find invariant, conserved amino acids in the central helix (Fig. 1b), the most conserved region of the protein. Third, a highly diverged ortholog from *D. mojavensis* in the *Drosophila* subgenus fully restores fertility to *D. melanogaster gdrd* mutants (Fig. 4a). Fourth, regardless of their abilities to rescue *gdrd* loss of function male sterility, most exogenously expressed Gdrd orthologs interact with axonemes and the insect ring centriole, indicating a conservation of functional protein-protein interactions. Altogether, these data suggest that the ancestral *gdrd* gene already possessed a critical role in sperm production in the common ancestor of *D. melanogaster* and *D. mojavensis*, roughly 40-43 million years ago. Furthermore, this work extends previous studies emphasizing structural conservation in *de novo* genes (Schmitz et al. 2018; Dowling et al. 2020; Lange et al. 2021; Chen et al. 2024; Middendorf et al. 2024; Peng and Zhao 2024) by showing that functional conservation can be equally long-standing.

### Extant Gdrd orthologs may have distinct, lineage-specific features

A surprising finding of this study is that the ability of *gdrd* orthologs to function in *D. melanogaster* spermatogenesis does not align neatly with phylogenetic distance. Two orthologs from the *melanogaster* species group—*D. yakuba* and *D. ananassae*—show weak or no ability to complement *gdrd* null alleles, despite having higher sequence conservations (Fig. 4a). By contrast, an ortholog from the *Drosophila* subgenus can fully rescue *gdrd* null fertility defects (Fig. 4a). Our localization analyses suggest that the ability to interact with the axoneme is one key determinant of rescue, as orthologs that strongly localize to this structure can substitute for the native gene (Fig. 4a, 5a). Furthermore, these observations underscore that Gdrd functions at this structure. Indeed, structure-function analyses of the *D. melanogaster* protein indicate that its interactions with the axoneme and insect ring centriole are mediated by the central helix, which also tends to be the most conserved region of Gdrd (Figure S3). A simple model to explain variability in rescue ability may be that amino acid substitutions in the central helix can either strengthen or weaken interactions. Such a mechanism would require constant co-evolution with a binding partner to maintain interactions (Lukeš et al. 2011; Hochberg et al. 2020; Chen et al. 2024).

Alternatively, the N- and C-terminal IDRs may also be critical determinants in the ability of a *gdrd* ortholog to substitute for the *D. melanogaster* protein. Across the genus, extant *gdrd* orthologs vary in their N- and C-terminal lengths, with some orthologs in the *repleta* and *melanica* species groups reaching close to twice the size of the *D. melanogaster* Gdrd. Interestingly, Gdrd orthologs in these two species groups also have aspartic acid/glycine rich regions, and their N-termini often carry a higher net negative charge. Indeed, such lineage specific changes underscore the potential importance of the N- and C-termini in modulating Gdrd function. Furthermore, although we cannot rule out protein instability, N- and C-terminal Gdrd truncation are incapable of rescuing *gdrd* mutant infertility to any degree despite the abilities of both truncated proteins to interact with the axoneme or the insect ring centriole (Fig. 3c,d). These observations suggest that the N- and C-termini of Gdrd orthologs may mediate lineage-specific contributions to Gdrd function.

Because the N- and C-terminal regions are intrinsically disordered, their amino acid sequences appear less conserved, especially when using traditional homology detection methods like BLASTP. In this study, we used k-mer-based SHARK analysis, SHARK-Dive (Chow et al. 2024) and SHARK-Capture (Chow et al. 2025), to identify conservation based on physicochemical properties. Analysis of orthologs across the *Drosophila* subgenus indicates that the C-terminus of the rescuing *D. mojavensis* ortholog shows a high level of physicochemical similarity with termini in *melanogaster* species group orthologs. Furthermore, this analysis revealed several shared physicochemical motifs within both IDRs of rescuing orthologs (*D. mojavensis*, *melanogaster* group) that are absent in *D. virilis*, potentially explaining the *D. virilis* protein’s weaker axonemal localization (Fig. 5,6). These motifs could have been present within the genus ancestor and then weakened or lost in various sublineages. Alternatively, these motifs and physicochemical properties might have arisen in *melanogaster* species group orthologs and convergently in the *D. mojavensis* ortholog. The latter scenario is consistent with the SHARK-Dive score for the *D. mojavensis* C-terminal IDR being a high outlier amongst *Drosophila* subgroup orthologs (Fig. S14).

At present, it is unknown if these changes in size, physicochemical properties, or motifs occur in response to changes in testes physiology, to alterations in reproductive programs, or if they are generally inconsequential. Our work, however, suggests that Gdrd orthologs have differences in intrinsic protein qualities, which may be important to their ability to function. Interestingly, the non-rescuing *D. ananassae* ortholog, despite being more closely related to the *melanogaster* group than *D. mojavensis* and containing the same conserved terminal motifs as rescuing species (Fig. 6), shows the highest instability during MD simulations (Fig. 7), suggesting that excessive structural lability could potentially impair its ability to complement the loss of the native gene. By contrast, the distant *D. mojavensis* ortholog has both the most stable structure of all tested orthologs—including *D. melanogaster*—and also the full capacity to substitute for the *D. melanogaster* protein. Interestingly, western blot analysis indicates that *D. ananassae* and *D. grimshawi* Gdrd proteins, both of which exhibit instability in MD simulations, are poorly expressed in a *D. melanogaster* testes, while the *D. mojavesis* protein shows the highest expression (Fig. 3a). This potentially underscores that amino acid sequence changes have led to changes to intrinsic properties of these proteins. A future avenue of inquiry may explore the relationship between physicochemical changes in termini and intrinsic protein qualities as predicted by MD simulations. Together with the SHARK analysis, these observations provide correlative support for a model in which lineage-specific changes in the termini contribute to functional divergence, a prediction that could be directly tested by future domain-swap experiments.

In our ortholog gene swap experiments, we also observed distinct subcellular localization patterns among several Gdrd orthologs. These novel patterns could stem either from lineage-specific evolution leading to functional innovation in their native species or from mislocalization in *D. melanogaster* caused by weakened interactions with host species proteins. However, we favor the former explanation for the *D. mojavensis*, *D. virilis*, and *D. ananassae* orthologs, which have stable and distinctive subcellular localizations during spermatid elongation. Interestingly, expanded Gdrd localization frequently involves membranous structures, such as the nuclear envelope in *D. mojavensis* and *D. virilis*, the acroblast/ Golgi in *D. ananassae*, and the plasma membrane in *D. grimshawi*. The *D. yakuba* protein also associates with an unidentified spherical organelle during elongation. This pattern may reflect an intrinsic membrane affinity of Gdrd, consistent with reports showing that intrinsically disordered proteins often mediate membrane binding (MacAinsh et al. 2025). In the context of *de novo* gene evolution, rapid sequence and physicochemical shifts within IDRs may drive divergence in Gdrd’s membrane or organellar associations, which vary in their chemical and physical properties. Supporting this idea, studies in yeast suggest that membranes can act as evolutionary incubators for *de novo* evolved gene products by providing stable environments that prevent protein aggregation or purge while allowing for the refinements of protein-protein interactions (Vakirlis et al. 2020; Houghton et al. 2024). Indeed, two other *Drosophila* orphan genes, *saturn* and *katherine johnson*, also encode membrane-associated proteins essential for male fertility (Gubala et al. 2017; Guay et al. 2025), suggesting that this evolutionary strategy may be commonly used in *Drosophila*. Interestingly, *goddard* orthologs that restore *D. melanogaster* fertility show broad localization to tubulin-associated, membrane-proximal structures such as the axoneme, ring centriole, and spermatid nuclear envelope, raising the possibility that the ancestral Gdrd could have functioned at membrane-linked microtubular interfaces. Whether nuclear cap localization represents an ancestral feature or a lineage-specific expansion in the *repleta* and *virilis* species groups, however, remains unresolved.

### Conservation and structural maintenance over a deep evolutionary timescale along with divergence

Collectively, our data are consistent with an evolutionary history in which core, rescue-relevant properties of Gdrd (e.g., helix stability, axoneme and insect ring centriole interaction) have been maintained over deep timescales, while other features diverged (Fig. 8). This pattern may be broadly representative of orphan genes that experience rapid sequence evolution and contain IDRs, such as genes encoding *de novo* originated or rapidly evolving, disordered proteins. Just as the gain and loss of *de novo* genes occur along a continuum (Carvunis et al. 2012; Heames et al. 2020; Iyengar and Bornberg-Bauer 2023), their functional trajectories after emergence likely follow a similar path, shaped by the binding promiscuity and polyvalency inherent to their IDRs. Such flexibility may promote coevolution with diverse interaction partners across lineages, enabling rapid adaptation for alternative purposes (Bornberg-Bauer et al. 2021; Jemth 2025).

**Figure 8:**
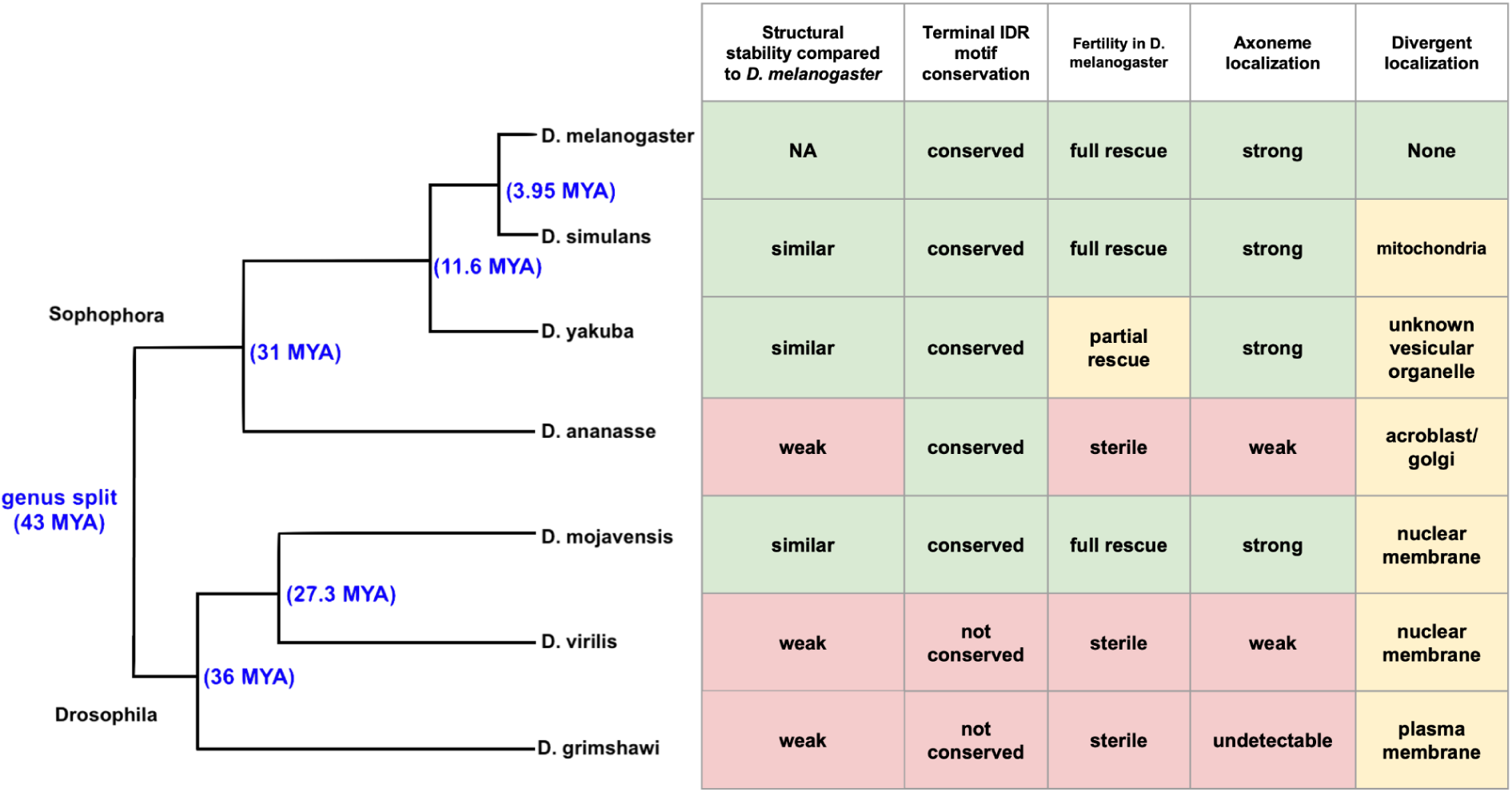
Summary of structural, k-mer, and gene swap analyses. Orthologs that lack function in gene swap assays typically exhibit weaker structural stability in MD simulations, have lost conserved terminal IDR motifs, or both. Most orthologs also exhibited divergent localization patterns. Divergence times (in blue brackets and font) were estimated using TimeTree. NA = Not Applicable.

Our experiments also suggest that amino acid sequence divergence in orphan genes can have major functional consequences within a gene swap context, as exogenously expressed orthologs often show diminished or lost activity and altered subcellular localizations. While structural conservation commonly suggests functional conservation, a case somewhat comparable to *gdrd* is seen with *bag of marbles* (*bam*), a well-studied orphan gene essential for *D. melanogaster* male and female fertility (McKearin and Spradling 1990). Like Gdrd, Bam lacks identifiable binding domains and contains intrinsically disordered regions (Lamb et al. 2020; Bubnell et al. 2022; Farfán-Pira et al. 2023; Baker et al. 2024). However, CRISPR/Cas9 knockouts in multiple *Drosophila* species have shown that *bam*’s essential role in fertility has been independently lost in several *melanogaster* group lineages (Lamb et al. 2020; Bubnell et al. 2022; Farfán-Pira et al. 2023; Baker et al. 2024), even though the gene has an equivalent function to *D. melanogaster* in a divergent member of the *Drosophila* subgenus, *D. americana* (Arnce et al. 2025). This suggests that lineage restricted genes conserved over a deep evolutionary timescale can lose (and possibly re-acquire) essential roles in fertility.

While our experiments suggest that amino acid sequence divergence can lead to diminished or lost function in a gene swap context, this approach has inherent limitations that must be considered. One potential shortcoming is that rescue constructs may not achieve expression levels equivalent to the endogenous locus. While RT-PCR indicates that all ortholog expression constructs produce equivalent or near equivalent levels of transcript (Fig. S4), protein abundance as determined by western blot analysis indicates that two orthologs—*D. ananassae* and *D. grimshawi*—are detected at lower levels in comparison to the *D. melanogaster* protein (Fig. 3a). As MD simulations suggest potential instabilities for these orthologs (Fig. 7), it is possible that lower expression may be a feature intrinsic to the amino acid sequences of these proteins. Furthermore, differences in an ortholog’s size, structures, and physiochemical properties (e.g., charge) likely contribute to the observed differences in western blot analysis, which can further confound interpretation. The level of protein produced from each orthologous gene could potentially be boosted by integration into the endogenous *D. melanogaster gdrd* locus (Adikusuma et al. 2017; Saint-Leandre et al. 2020; Brand and Levine 2022), though this would not change the orthologs’ inherent physicochemical properties. Boosting expression might also exacerbate spermatid elongation defects for orthologs (e.g., *D. ananassae*) that interfere with *D. melanogaster* spermatogenesis, further complicating interpretation.

A second limitation arises from the exogenous nature of gene swap assays. Although testis architecture and function are broadly conserved across *Drosophila*, interspecies physiological differences persist (Schärer et al. 2008; Lüpold et al. 2016; Alpern et al. 2019). Addressing this limitation will require direct examination of orthologous Gdrd proteins in their native testes using CRISPR-generated knockouts in combination with either specific antibodies or HA-tagged knock-ins in non-*melanogaster* species (Lamb et al. 2020; Bubnell et al. 2022; Farfán-Pira et al. 2023; Baker et al. 2024; Arnce et al. 2025). Such experiments will also illuminate whether the divergent subcellular localization patterns we observed of many orthologs when expressed in *D. melanogaster* match conspecific patterns.

In conclusion, our findings suggest that the common ancestor of the *Sophophora* and *Drosophila* subgenera likely expressed a Gdrd protein that already possessed the core interactions and activities required for spermatogenesis in present-day *D. melanogaster*. The conservation of transition zone localization and the association of axonemal localization with function underscore that proper Gdrd positioning at these structures is critical for its role. However, our gene swap analyses also indicate that Gdrd function might have diverged across lineages. Orthologs from *D. mojavensis* and *D. virilis* show distinct and expanded localization patterns in *D. melanogaster*, suggesting the possibility that they could have broader subcellular roles in their native species, while the conserved *D. ananassae* ortholog may have evolved new interaction preferences. Future studies will determine whether these altered localization patterns correspond to expanded or novel gene functions.

## MATERIALS AND METHODS

### Fly stocks and husbandry

For a complete list of fly strains used in this study, please see the Reagents Table. For all experiments, flies were raised on standard cornmeal molasses media (Guay et al. 2025) at 25°C without a light-dark cycle.

### Sequence alignment and truncated *D. melanogaster* Gdrd design

Based on our earlier study, which included concordant experimental results from circular dichroism spectroscopy and 2D NMR spectroscopy (Lange et al. 2021), we subdivided *D. melanogaster* Gdrd protein into three regions: a 40 amino acid (aa) disordered N-terminus, a 38 aa central helix, and a 35 aa disordered C-terminus. Homology analyses between the *D. melanogaster* protein and *Drosophila* genus orthologs were performed using pairwise BLASTP (Altschul et al. 1997). BLASTP was run locally (BLAST+ 2.16.0+) in an all-against-all manner using the short-query preset (-task blastp-short) with an E-value threshold of 0.05; all other parameters were defaults for blastp-short. Output was parsed in tabular format (-outfmt “6 qseqid sseqid nident length bitscore”), and for each query–subject pair the best High-Scoring Pair (HSP) by bitscore was retained. Percent identity was computed as nident/alignment-length × 100. As distant orthologs within the *Drosophila* subgenus were highly divergent at the N- and C-termini of the protein, amino acid sequence conservation (percent identity) was alternatively determined within the context of a Clustal Omega (ClustalO) multiple sequence alignment (Sievers et al. 2011) and analyzed using ggmsa and ggplot2 in R v4.2.2 (Wickham 2011; Zhou et al. 2022). Truncated versions of *D. melanogaster* Gdrd were designed based on AlphaFold2 (Jumper et al. 2021) predictions, experimental structure data (Lange et al. 2021), and alignment of the central helix (Fig. 1b). The N-terminal truncation (Gdrd ΔN) removes 36 amino acids following the initial methionine. The C-terminal truncation (Gdrd ΔC) removes the final 35 amino acids of the protein, immediately downstream of the central helix. We used the DeNoFo toolkit (Dohmen et al. 2025) to annotate the methods used to identify Gdrd in earlier studies (Gubala et al. 2017; Lange et al. 2021). The DeNoFo annotation file available in the supplementary material.

### Alignment-free analysis of terminal IDRs

Based on the sequence alignment of the central helices (Fig. 1b), sequences of the disordered termini were extracted from each ortholog. Homology of each ortholog’s N-and C-termini to those of *D. melanogaster* Gdrd was calculated using the alignment-free, k-mer-based tool SHARK-Dive (Chow et al. 2024). Similarity was normalized with D. *melanogaster* Gdrd self-comparison set as 1. Shared motifs were identified using SHARK-Capture in default mode (Chow et al. 2025). Overall charge of tails was calculated using a custom python script (D, E = −1, K, R, H = +1, other amino acids = 0).

### Transgenic flies

All transgenic constructs used in this analysis constitute modifications of the previously reported *gdrd:HA* rescue (Lange et al. 2021). For ortholog gene swaps, the *D. melanogaster*, *D. simulans* (LOC6738055), *D. yakuba* (LOC6534118), *D. ananassae* (LOC6507245), *D. mojavensis*, *D. virilis*, and *D. grimshawi gdrd* coding sequences were first codon-optimized using GENEius software (Eurofins MWG Operon). Amino acid sequences for *D. mojavensis*, *D. virilis*, and *D. grimshawi* Gdrd orthologs were previously documented (Gubala et al. 2017). Forty-nucleotide upstream and downstream homology arm sequences (corresponding to the 5’UTR and HA-tag sequences in the original rescue, respectively) were then attached to each codon-optimized gene sequence prior to synthesis (Eurofins MWG Operon). The upstream regulatory regions containing the *D. melanogaster gdrd* 5’UTR and the downstream regulatory regions containing the Hemagluttanin (HA) tag were PCR amplified using Q5 High Fidelity Polymerase (NEB). These PCR fragments, along with the synthesized codon-optimized gene, were cloned into a XbaI/AscI-linearized w+attB plasmid (Sekelsky, Addgene plasmid 30326) using Gibson Assembly (NEB). Gdrd ΔN and Gdrd ΔC constructs are modifications of the codon-optimized *mel gdrd* rescue construct. Sequences that omit these protein regions were amplified using Q5 High Fidelity Polymerase (NEB) and extended primers designed with 20 bp homology regions. These DNA fragments were then cloned into w+attB plasmid using Gibson assembly. To control for chromatin position effects on expression, each rescue construct was then phiC31 integrated into the *PBac{y^+^-attP-9A}VK00020* (BL24867) docking site (Rainbow Transgenics). Upon establishing transgenic lines in the *D. melanogaster gdrd* null background, we used PCR and sequencing to re-confirm ortholog insertions and the absence of the endogenous gene. To maximize gene expression from transgenes, all tested males carry two copies of each rescue insertion, unless otherwise noted. RT-PCR was used to evaluate transcript levels in sexually mature males of carrying each transgene in the *gdrd* null background. Primers located in the 5’ and 3’ UTRs of *gdrd* were used to amplify the full length protein-coding sequence (including the 3xHA tag) of each transgene from cDNA as previously described (Rivard et al. 2021). As a control for cDNA synthesis and to detect any contaminating genomic DNA, the housekeeping control gene *RpL19* was also amplified from each cDNA sample. See the Reagents Table for primer and synthesized DNA sequences.

### Western blot analysis

10 testes pairs were dissected in PBS buffer and then transferred to 45 uL Ripa lysis buffer (Thermo Scientific Catalog No. #89900). 15 uL of 4X sample buffer (200 mM Tris pH 6.8, 8% Sodium dodecyl sulfate (W/V), 20% glycerol, 0.1% bromophenol blue, 10% β-mercaptoethanol) was added prior to homogenization and sample boiling. An equivalent of 4 testes were run in each lane of either a 15 or 20% SDS-PAGE gel. Resolved proteins were then transferred to PVDF membrane (pore size 0.2 μm; BioRad) using wet transfer (Towbin buffer) prior to standard western blotting immunodetection. Blots were blocked in 2% milk and probed with rabbit anti-HA (Cell signaling; 1:1000) and mouse anti-actin (MP Biomedicals/VWR; 1:4000). Primary antibodies were detected with HRP conjugated secondaries (1:5000). HRP activity was detected using Pierce ECL 2 Western Blotting Substrate (Thermo Scientific). See Reagents Table for details about antibodies.

### Fertility assay

To assess the functional abilities of *gdrd* orthologs and truncations in *D. melanogaster*, we performed single male fertility assays by mating individual males with two *Canton-S* virgin females. All males were collected as virgins and then aged in uncrowded conditions for four to six days prior to mating. The age range of virgin females fell between four to eight days. Likewise, virgin females used in the assay were isolated in uncrowded condition and reared on a high yeast diet prior to mating to increase fecundity. Mating and egg laying proceeded for 48 hours post-crossing before parents were discarded. Progeny number was determined by counting the number of pupal cases on the side of each vial 10 days (∼216 hours) after setting the cross. Final sample sizes for each genotype were *n* = 25-30 males, depending on the experiment. Progeny count data were analyzed in Microsoft Excel using *t*-tests with unequal variances.

### Fluorescent labeling and confocal microscopy of *Drosophila* whole testes and testes squashes

Antibody labeling of whole testis and testis squashes were performed as previously described in Lange *et al*. (2021) and Sitaram *et al*. (2014), respectively. In brief, processed tissues were fixed following dissection in 4% paraformaldehyde/ 1X phosphate buffered saline (PBS) for 20 minutes. After washing the fixative away, tissues were blocked for 1-2 hours in 1X PBS containing 3% bovine serum albumin, 5% normal goat serum, and 0.1% Triton-X. The samples were then incubated overnight in rabbit anti:HA (C29F4, Cell Signaling Technologies) diluted at 1:100 in blocking solution. The following day, the antibody was washed off before applying goat anti-rabbit Alexa Fluor 488 conjugated secondary (A-11008, ThermoFisher Scientific) diluted at 1:200 in blocking solution. The secondary was then washed away prior to mounting samples in Vectashield Plus Antifade Mounting Medium Plus DAPI (Vector labs). To analyze individualization complexes during spermatogenesis, fixed whole testes were incubated overnight with TRITC conjugated phalloidin (R415, Life Technologies/ Molecular Probes) diluted at 1:200 in PBS containing 0.1% Triton-X. Microscopy was performed on an SP8 X confocal microscope (Leica Microsystems) using HC PL APO CS2 20x/0.75 ILL and HC PL APO CS2 63x/1.40 oil objectives. DAPI, Alexa Fluor 488, and TRITC were excited using 405, 488, and 546 wavelengths, respectively. Post-acquisition processing was performed using ImageJ Fiji (version 1.0) (Schindelin et al. 2012).

### Structure predictions and MD simulations

Structure predictions were performed with AlphaFold2 (v.2.3.1; Database cut-off date: 2022-12-19) (Jumper et al. 2021). MD simulations were performed using GROMACS 2021.2 (Páll et al. 2020) on AlphaFold2 predictions with highest mean predicted local distance difference test (pLDDT) score. Structures were prepared using the OPLS-AA/L force field (Shivakumar et al. 2010) and solvent SOL. Structures were solvated in a cubic box of SPC/E water with 10-Å clearance and the electrostatic charge neutralized by the addition of sodium atoms, followed by energy minimization and equilibration. Triplicate 250 ns simulations were run in an NPT ensemble using a V-rescale modified Berendsen thermostat at a temperature of 300 K and a Parrinello-Rahman barostat at a pressure of 1 atm, periodic boundary conditions, and a particle mesh Ewald summation with a grid spacing of 1.6 Å and fourth order interpolation. Custom python and bash scripts were used for analysis of MD simulations (Backbone RMSD, α-C RMSF, radius of gyration). For each species, triplicate 250 ns MD simulation snapshots were aligned using PyMOL v3.0.5 (Schrödinger 2015). Structures were visualized using PyMOL v3.0.5 and ChimeraX-1.7.1 (Pettersen et al. 2021). Statistical analyses and plotting were performed using Python v.3.10.13 (libraries: numpy (Harris et al. 2020), pandas (Mckinney 2011), scipy (Virtanen et al. 2020), scikit-posthocs (Terpilowski 2019), matplotlib (Hunter 2007)). For RMSD and radius of gyration, we analyzed equilibrated trajectories 50-250 ns subsampled at 100-frame intervals, yielding ∼600 observations per species (200 timepoints × 3 replicates). For RMSF, all residues within each region (N-terminus, core, C-terminus) across three replicates were analyzed (n=100-300 per species). Kruskal-Wallis H-tests assessed overall differences, followed by Dunn’s post-hoc tests with Bonferroni correction (α=0.05, 21 comparisons). NaN values were excluded.

### Use of Artificial Intelligence

After completing data collection and analysis and drafting and editing the full manuscript, we used GPT-5 to condense the text. We then manually edited prior to submission. All revisions were done with manual editing.

## Supporting information

Reagents table

Raw data file

## Acknowledgements

We thank Dr. Tomer Avidor-Reiss (Department of Biological Sciences, University of Toledo, Toledo, OH) for providing us with the *unc:EGFP* transgenic fly strain. We thank Dr. Emily Rivard and four reviewers for providing helpful feedback on the manuscript. P.H.P, K.L.M and G.D.F were supported by NSF RUI grant (2212972) awarded to G.D.F. E.B.B. and A.L. received funding from Volkswagen foundation grant code 98183. E. B. B. was supported by HFSP (Human Frontiers of Science Programme, RGP0006/2013 and RGP0006/2013) and the DFG (Deutsche Forschungsgemeinschaft BO-2544/20-1;503272152). L.A.E. has been supported by EMBO Scientific Exchange Grant 10944. Structure predictions and MD simulations were performed on the HPC cluster PALMA II of the University of Muenster, subsidised by the DFG (INST 211/667-1).

## Data Availability Statement

Scripts for sequence and structural analysis are deposited on GitHub: https://github.com/ArsLeicholt/gdrd_analysis. Supplementary data can be found on Zenodo: https://doi.org/10.5281/zenodo.15173276. Data underlying figures from the manuscript is provided in supplemental material.

