## Supplemental figures and text for "Orthologs of an essential orphan gene vary in their capacities for function and subcellular localization in *Drosophila melanogaster*"

By Prajal H. Patel et al.

### Supplementary Text:

#### Analysis and interpretation of Gdrd and Unc localization patterns during spermatogenesis

Our analysis of codon-optimized *D. melanogaster* Gdrd expression and localization in isolated cysts recapitulates previous whole-mount findings (Lange et al. 2021) (Figure S5). Gdrd expression initiates in mature spermatocytes, where the protein exhibits diffuse cytoplasmic localization (Figure S5b). At this stage, spermatocytes contain four Unc-positive basal bodies (Figure S5b) that anchor cilia within a plasma membrane cap (Baker et al. 2004; Riparbelli et al. 2012). Besides acetylated tubulin and basal body components (Basiri et al. 2014), these structures also contain components of the transition zone/ ciliary gate, including MKS1, Cep290, BD91/ BD92, and Cby (Pratt et al. 2016). Gdrd shows no enrichment at these ciliary structures, suggesting that the protein does not interact with any of the aforementioned components. Therefore, the ciliary gate and basal bodies of elongating spermatids are either compositionally distinct or are post-translationally modified during spermatid elongation to recruit Gdrd to this structure.

During spermatid elongation initiation, Gdrd exhibits cytoplasmic localization (i.e., nuclear and mitochondrial exclusion) and colocalization with Unc at the basal body, transition zone, and axoneme (Figure S5c). As cyst elongation proceeds in early elongating spermatids, this pattern becomes restricted. Unc remains at the basal body and transition zone but no longer associates with the axoneme (Figures S5d). By contrast, Gdrd enriches distally to the Unc-positive basal body, to either a distinct basal body or axonemal subdomain (Figure S5d). At these stages, Gdrd is both cytoplasmic and enriched at the axoneme. In late elongating spermatids, Gdrd expression is detectable but diminished (Figure S5d). In late elongating spermatid cysts, Gdrd colocalizes with Unc at the transition zone (Figure S5e), a structure that persists until the onset of sperm individualization (Vieillard et al. 2016). Our immunohistochemical analysis of Gdrd reveals that the protein disappears prior to this disassembly (Figure S5e).

In many insect species, the sperm axoneme is principally cytoplasmic, with only its distal tip sequestered within a membranous ciliary cap (Figure S5e) (Johnson 1922; Phillips 1970; Basiri et al. 2014). This compartmentalization is mediated by the insect ring centriole, an electron dense structure that lacks centriole-like organization (Phillips 1970). An analogous murine structure, the annulus, shares both protein components and functions with the insect ring centriole (Kwitny et al. 2010; Basiri et al. 2014; Hoque et al. 2024). In flies, this compartment is thought to promote efficient tubulin polymerization for axoneme elongation (Basiri et al. 2014). Although Gdrd decorates the axoneme throughout elongation, it never localizes beyond the transition zone, suggesting that the compartmentalized distal tip is either biochemically distinct from the cytoplasmic axoneme or that the ciliary gate bars Gdrd entry (Figure S5c, S5d). Altogether, these findings indicate that Gdrd associates with the cytoplasmic flagellum until a time point just before the start of spermatid individualization.

**Non-parametric statistical analysis of fertility and individualization complex data (Figure 4)**

In addition to the parametric statistical analyses presented in the main text, we also analyzed the fertility assay and individualization complex (IC) quantification data using non-parametric analyses. These analyses reached the same conclusions as the parametric tests. For the fertility assay data (Fig. 4a), a one-way ANOVA analysis revealed significant differences between male genotypes ($F_{9,290}$ = 579.1, $P < 10^{-178}$). Tukey-Kramer tests grouping all of the highly fertile genotypes in one group (all $P >$ 0.57), and the low-fertility *yakuba* ortholog and all non-rescuing genotypes into a second group (all $P > 0.95$). For the IC data (Fig. 4b), a one-way ANOVA ($F_{8,144}$ = 111.7, $P < 10^{-57}$) detected a significant difference across male genotypes. Pairwise Tukey-Kramer tests grouped the genotypes as follows: *sim* ($p < 0.002$ for all pairwise comparisons to other genotypes); $w^{1118}$, *mel*, *moj*; *yak* ($P < 0.0001$ for all pairwise comparisons to other genotypes); *ana, vir, gri*.

**(a)**

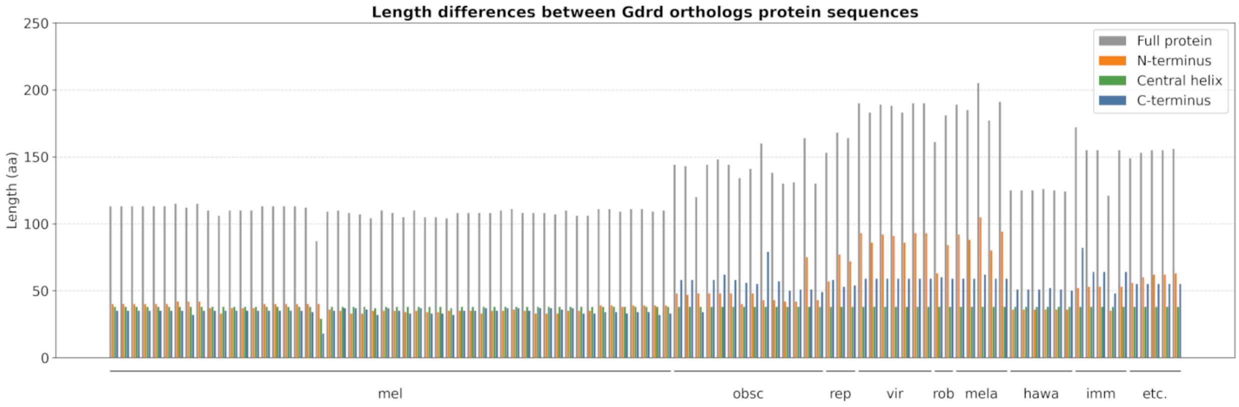


**(b)**

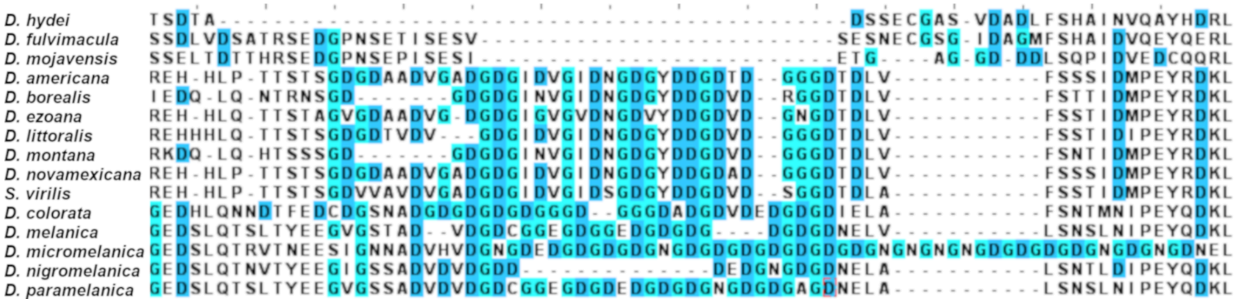


**Figure S1.** *Gdrd* orthologs across the *Drosophila* genus vary in overall sequence length. a) Among identified orthologs, central helix lengths are largely conserved. While total protein length within the *melanogaster* species group is generally similar, outgroup species tend to have modestly extended C-termini. In contrast, N-termini from the *repleta*, *virilis*, *robusta*, and *melanica* species groups within the *Drosophila* subgenus are typically longer than those of outgroup species. b) Clustal Omega alignments of representative Gdrd N-terminal regions from the *repleta*, *virilis*, and *melanica* species groups. The *virilis* and *melanica* groups contain glycine/aspartic acid–rich regions that are absent in the *repleta* group.

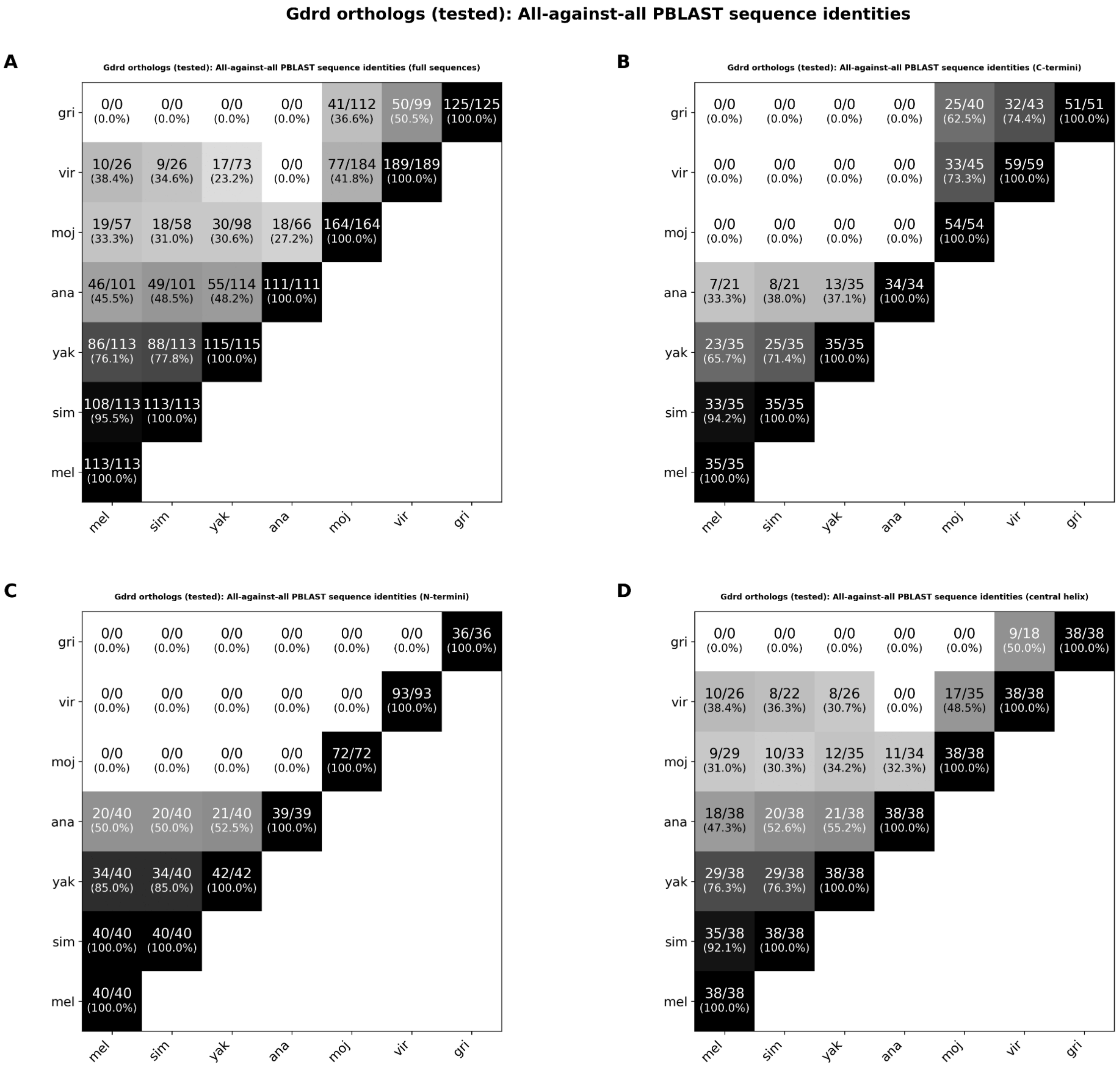


**Figure S2.** Pairwise BLASTP analysis of Gdrd orthologs. All possible pairwise comparisons were made between Gdrd orthologs from *D. melanogaster*, *D. simulans*, *D. yakuba*, *D. ananassae*, *D. mojavensis*, *D. virilis*, and *D. grimshawii*. The tables show the percent identity for comparisons of: (a) the full-length protein, (b) the N-terminus, (c) the C-terminus, and (d) the central helix region.

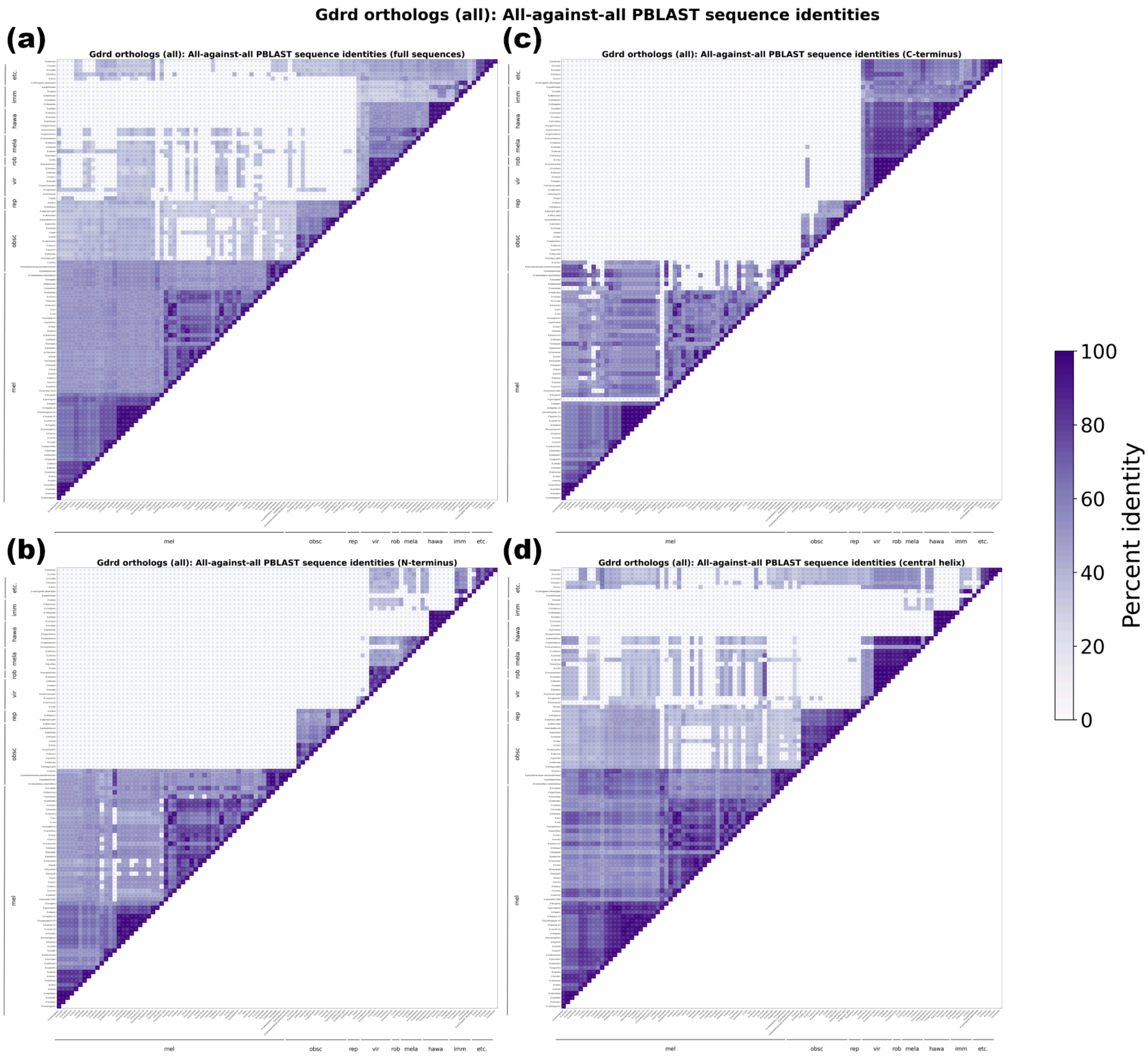


**Figure S3.** Pairwise BLASTP analysis of all identified *Gdrd* orthologs. The tables report percent identity for comparisons of a) the full-length proteins, b) N-termini, c) C-termini, and d) the central helix regions. Conservation is highest within the central helix. Among species of the *Drosophila* subgenus, C-termini are generally more conserved than N-termini, consistent with the greater variability in N-terminal sequence lengths among these species.

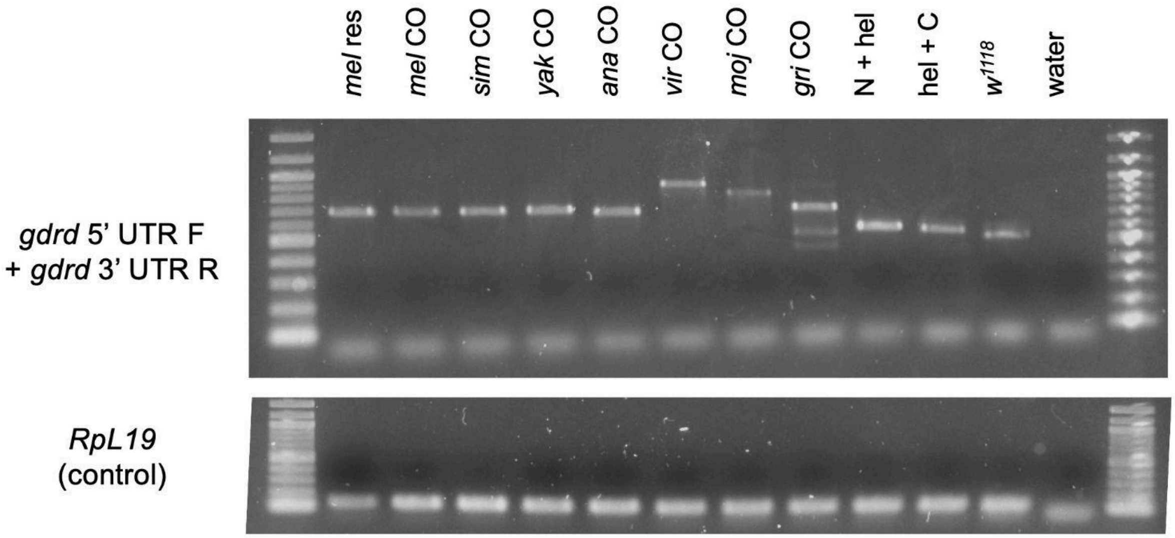


**Figure S4.** RT-PCR assessment of *gdrd* rescue construct transcript abundance. *Gdrd* transcripts were amplified from cDNA prepared from whole flies expressing rescue constructs in a *gdrd* null background, using primers targeting the shared *D. melanogaster* 5′ and 3′ UTRs, which are present in all constructs. Nearly all constructs had high levels of transcript abundance. Although transcript levels were slightly reduced for the *D. mojavensis* ortholog, expression is sufficient to fully rescue fertility in *D. melanogaster gdrd* null males (Fig. 4a).The *gdrd* product for the $w^{1118}$ strain is smaller than the *mel* rescue and *mel* CO lines due to the lack of the 3xHA tag. As a control for cDNA synthesis and to detect any potential genomic DNA contamination, the housekeeping gene *RpL19* was amplified using intron-spanning primers. No larger, intron-containing product was observed in samples. The smaller product in the water lane for *RpL19* corresponds to primer dimers.

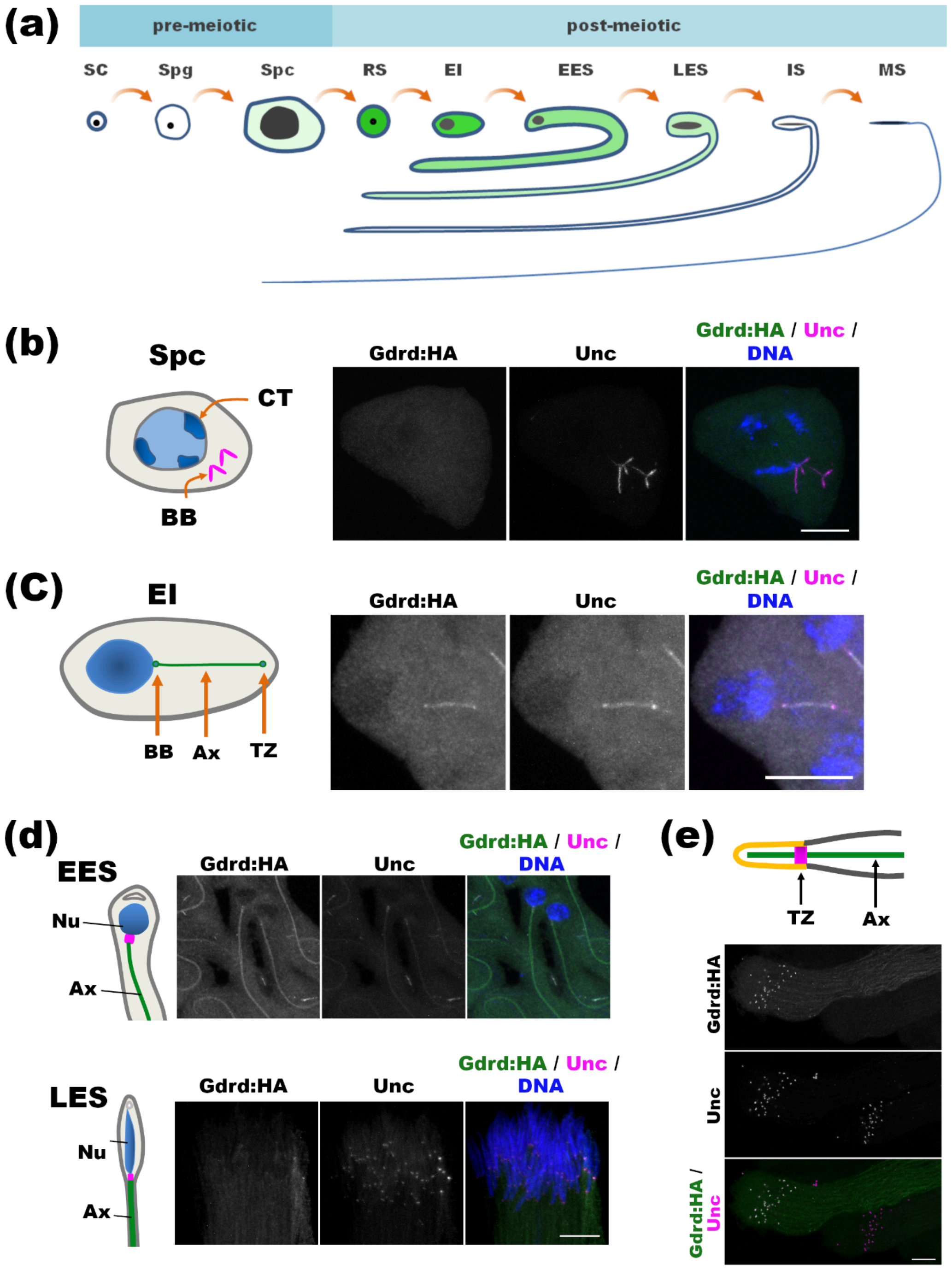


**Figure S5.** Codon-optimized *D. melanogaster* Gdrd gene swap recapitulates the expression and localization patterns of the wild-type rescue. a) Spermatogenesis progresses through a series of pre- and post-meiotic stages, including germline stem cells (SCs), spermatogonia (Spg), spermatocytes (Spc), round spermatids (RS), elongation initiation (EI), early and late elongating spermatids (EES and LES), individualizing spermatids (IS), and mature sperm (MS). Darker and lighter green shading, respectively, denote Gdrd's stage-specific higher or lower expression levels as assessed by immunofluorescence. b) Codon-optimized *D. melanogaster* Gdrd localizes throughout the spermatocyte cytoplasm but shows no enrichment at cilia-like basal bodies. c) *D. mel* Gdrd colocalizes with Unc at the basal body, migrating insect ring centriole, and the axoneme in spermatids undergoing elongation initiation. d) *D. mel* Gdrd decorates axonemes strongly in early elongating spermatid cysts and weakly in late elongating spermatids. e) At the distal end of elongated spermatid cysts, Gdrd co-localizes with Unc at the transition zone/ ciliary gates (top cyst). Both Gdrd expression and localization disappear prior to the disassembly of transition zones/ ciliary gates (bottom cyst). In each panel, accompanying schematic diagrams depict the stage-specific organization of spermatids and highlight the cellular structures shown in the corresponding micrographs (Nu, nucleus; CT, chromosome territory; Ax, axoneme; TZ, transition zone; BB, basal body). Genotype in all images: *unc:EGFP rescue, codon-optimized (CO) D. mel gdrd:HA rescue, Δgdrd*. All images are composed of confocal z-stacks. Scale bar for all images = 10 μm.

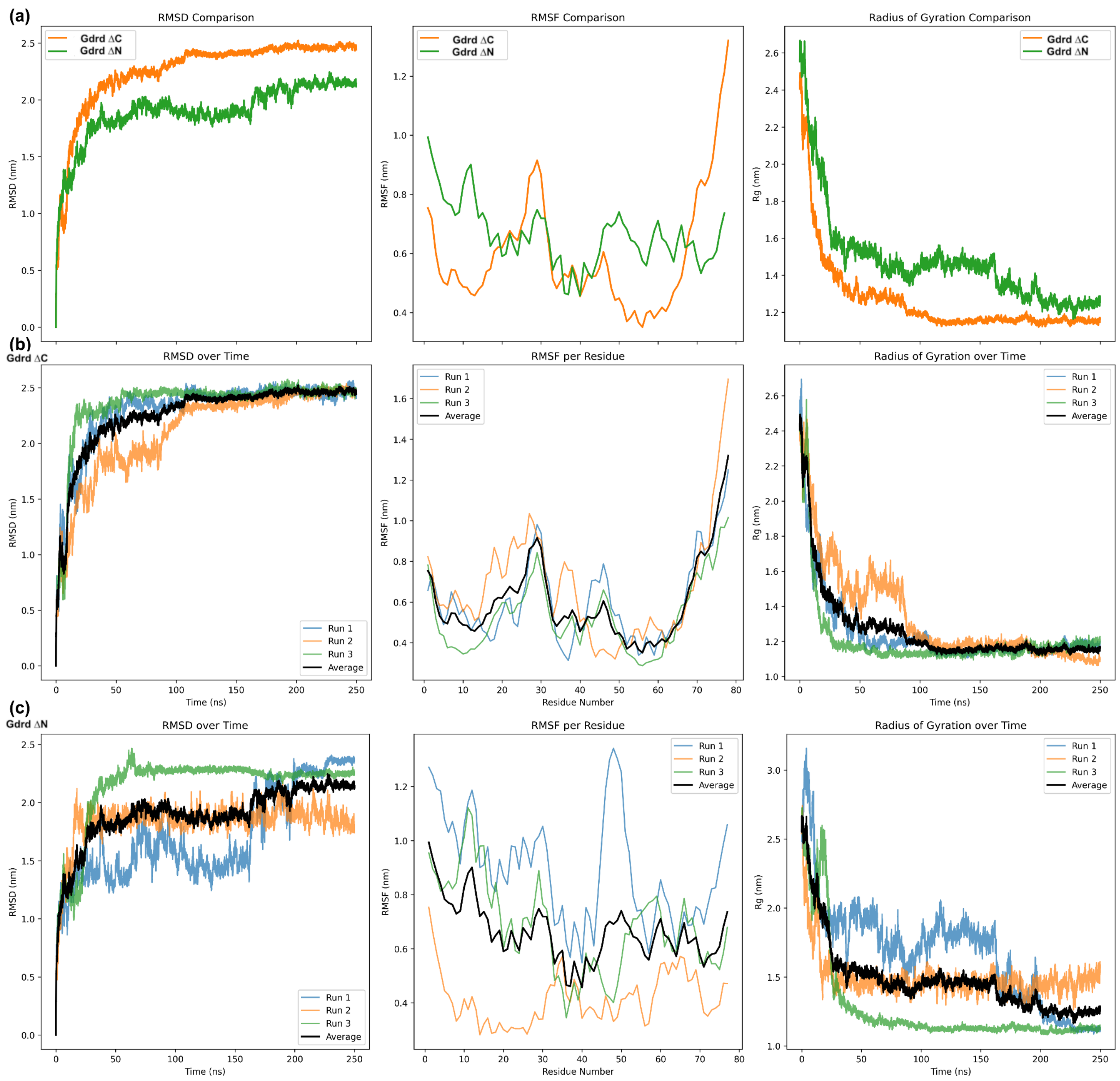


**Figure S6.** Comparison of molecular dynamics (MD) simulations to assess the stability of *D. melanogaster* Gdrd truncations. Left column shows backbone RMSD, center column shows $\alpha$-C RMSF, and right column shows radius of gyration. a) Comparison of average values for three simulations each of Gdrd ΔC and Gdrd ΔN truncations. b) Individual simulations (triplicate) for Gdrd ΔC truncation. c) Individual simulations (triplicate) for Gdrd ΔN truncation.

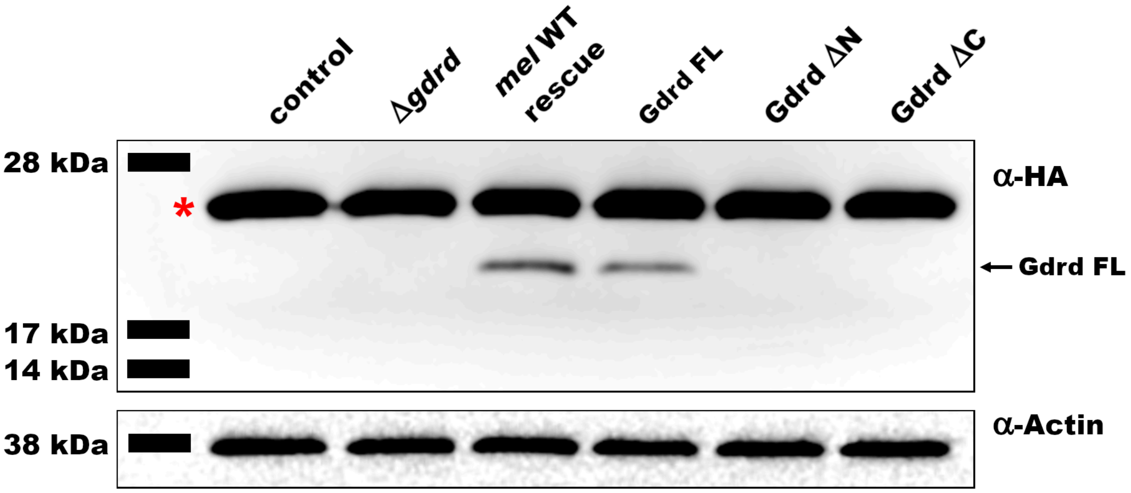


**Figure S7.** Western blot of protein lysates generated from *D. melanogaster* testes expressing HA-tagged full length or truncated Gdrd proteins. Truncated Gdrd proteins are difficult to detect on western blots. Control is *w*$^{1118}$ mutant genetic background. Red asterisk labels a non-specific band present in both control and experimental lysates. Actin serves as loading control.

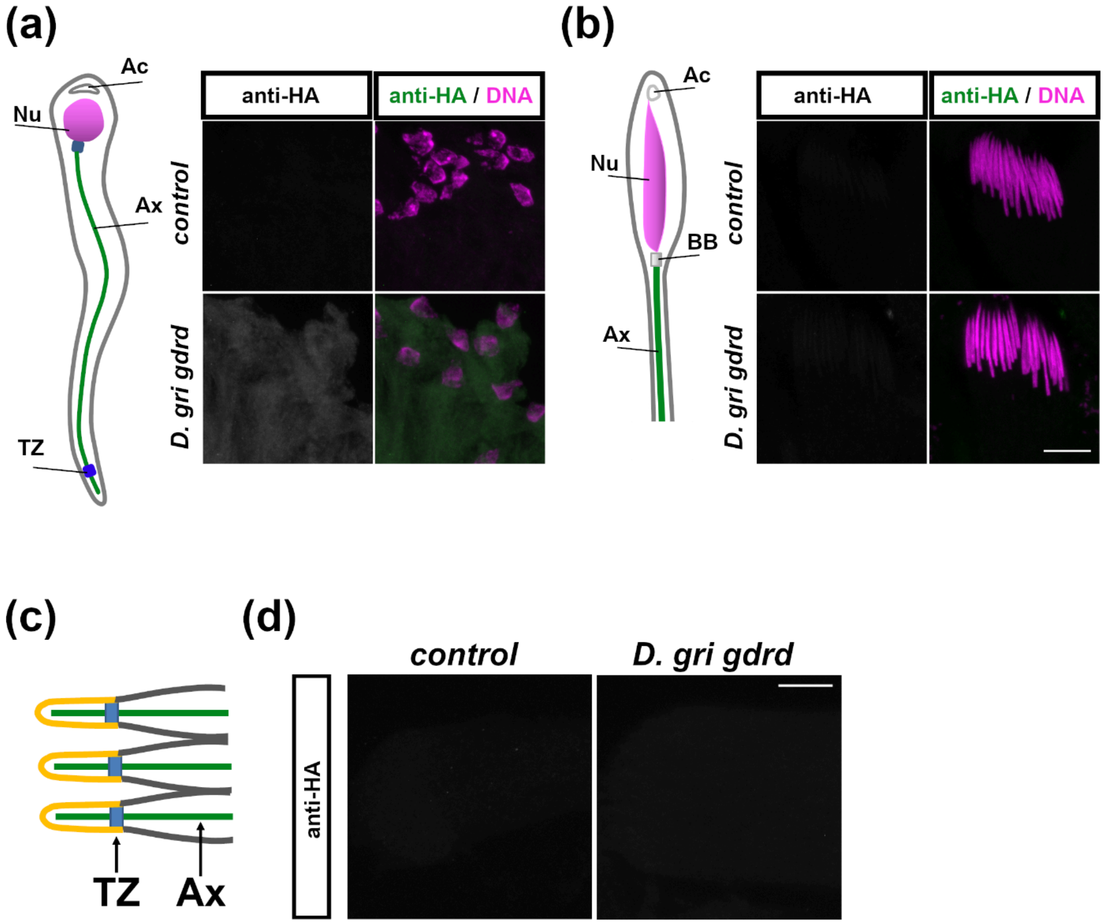


**Figure S8.** *D. grimshawi* Gdrd expressed in *D. melanogaster* testes fails to localize to either axonemes or transition zones. a) *D. grimshawi* Gdrd fails to localize to axonemes in early elongating spermatids. b) *D. grimshawi* Gdrd is undetectable in late elongating spermatids. c) *D. grimshawi* Gdrd is absent from transition zones at the distal ends of elongating cysts. In each panel, accompanying schematic diagrams depict the stage-specific organization of spermatids and highlight the cellular structures shown in the corresponding micrographs (Nu, nucleus; Ax, axoneme; TZ, transition zone; BB, basal body; Ac, acrosome). $w^{1118}$ served as the control in all panels. Scale bar = 10 μm for all images.

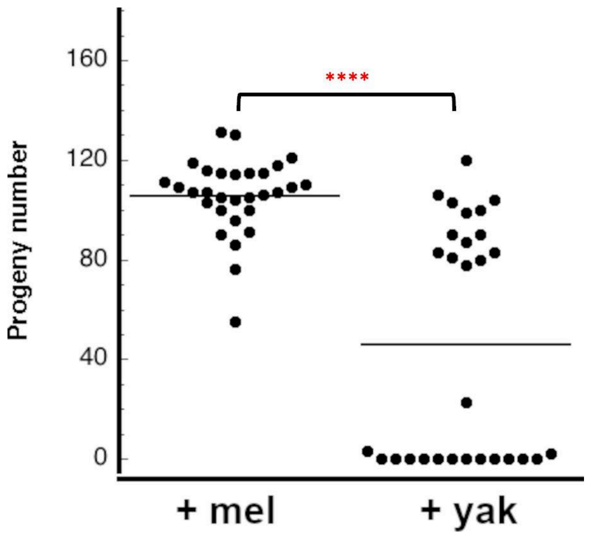


**Figure S9.** Replicate fertility assay of *D. yakuba* gene swap males. *D. yakuba* gene swap males exhibit bimodal fertility, with some males exhibiting near wild-type levels of fertility while other males are either sterile or near sterile. Statistical test: *t*-tests with unequal variance. **** = p < .0001

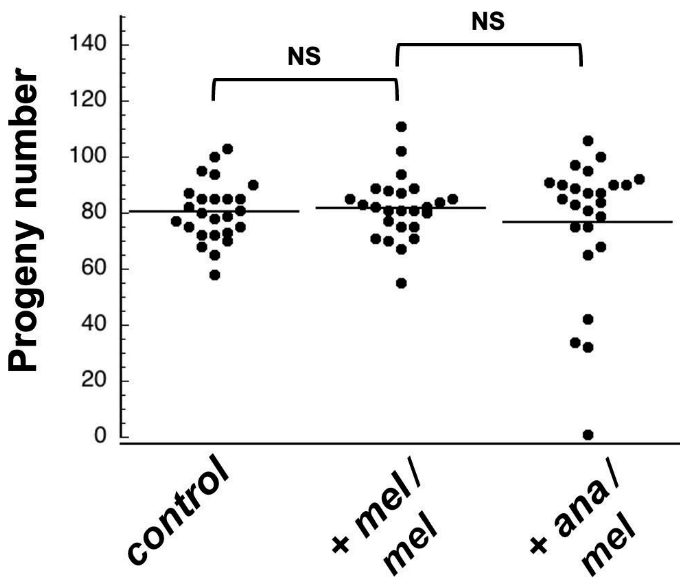


**Figure S10.** Spermatid elongation attenuation phenotypes in *D. ananassae* gene swaps are recessive. Transheterozygous male flies carrying one *D. ananassae gdrd* and one *D. melanogaster gdrd* rescue construct in a *gdrd* null background have wild type levels of fertility. Statistical test: *t*-tests with unequal variance. NS = Not Significant.

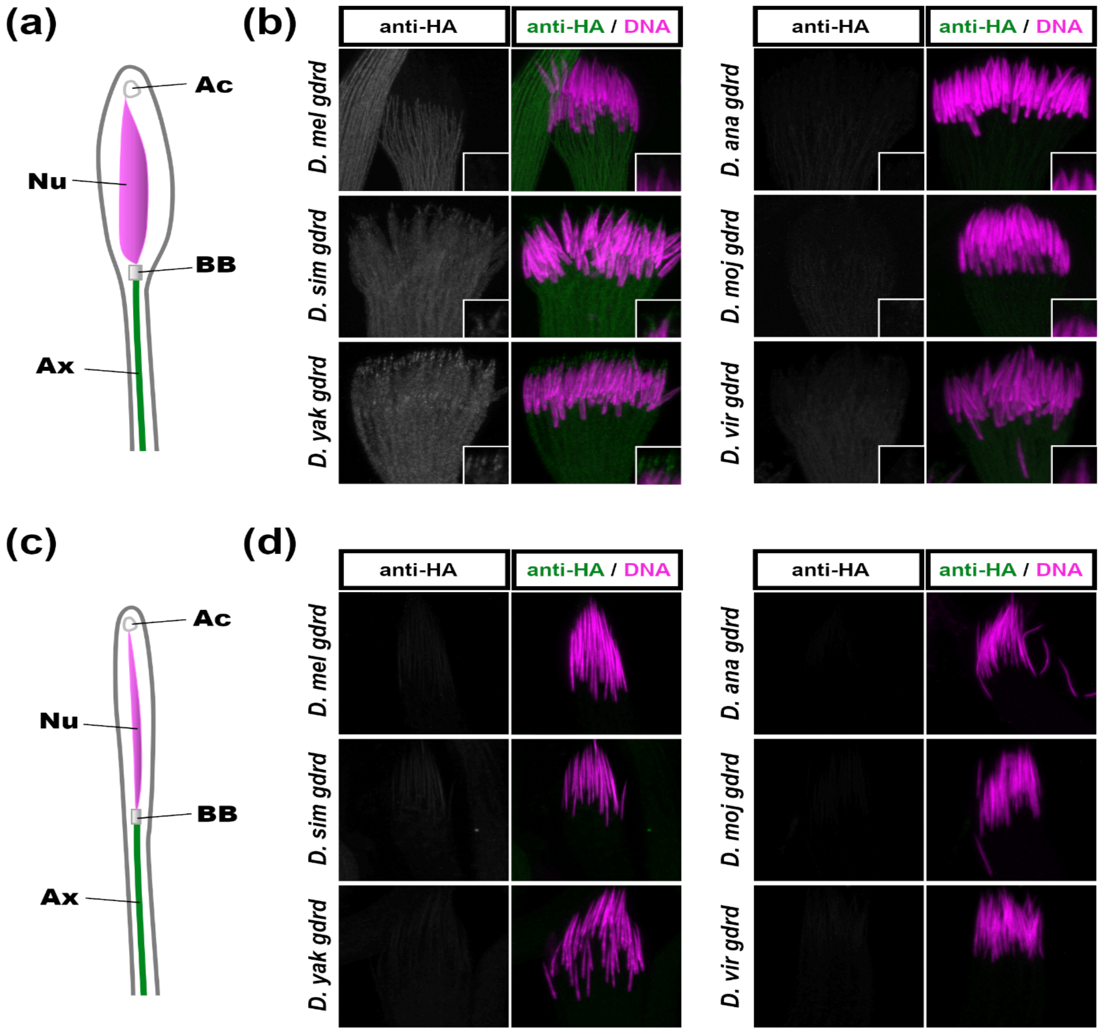


**Figure S11.** Localization of Gdrd orthologs in late elongating and individualizing spermatids. a-b) All Gdrd orthologs are detectable in late elongating spermatids. The *D. sim* and *D. yak* proteins appear enriched at the basal end of nuclei (see insets) near or at the acrosome, with the signal in *D. yak* being more robust. c-d) Gdrd expression is absent in all individualizing spermatids. In each panel, accompanying schematic diagrams depict the stage-specific organization of spermatids and highlight the cellular structures shown in the corresponding micrographs (Ac, acrosome; Nu, nucleus; BB, basal body; Ax, axoneme). Canoe or needle shaped nuclear morphology was used to stage spermatid cyst as either late elongating or individualizing spermatids, respectively. The basal end of each cyst is at the top of the image. Scale bar = 10 µm.

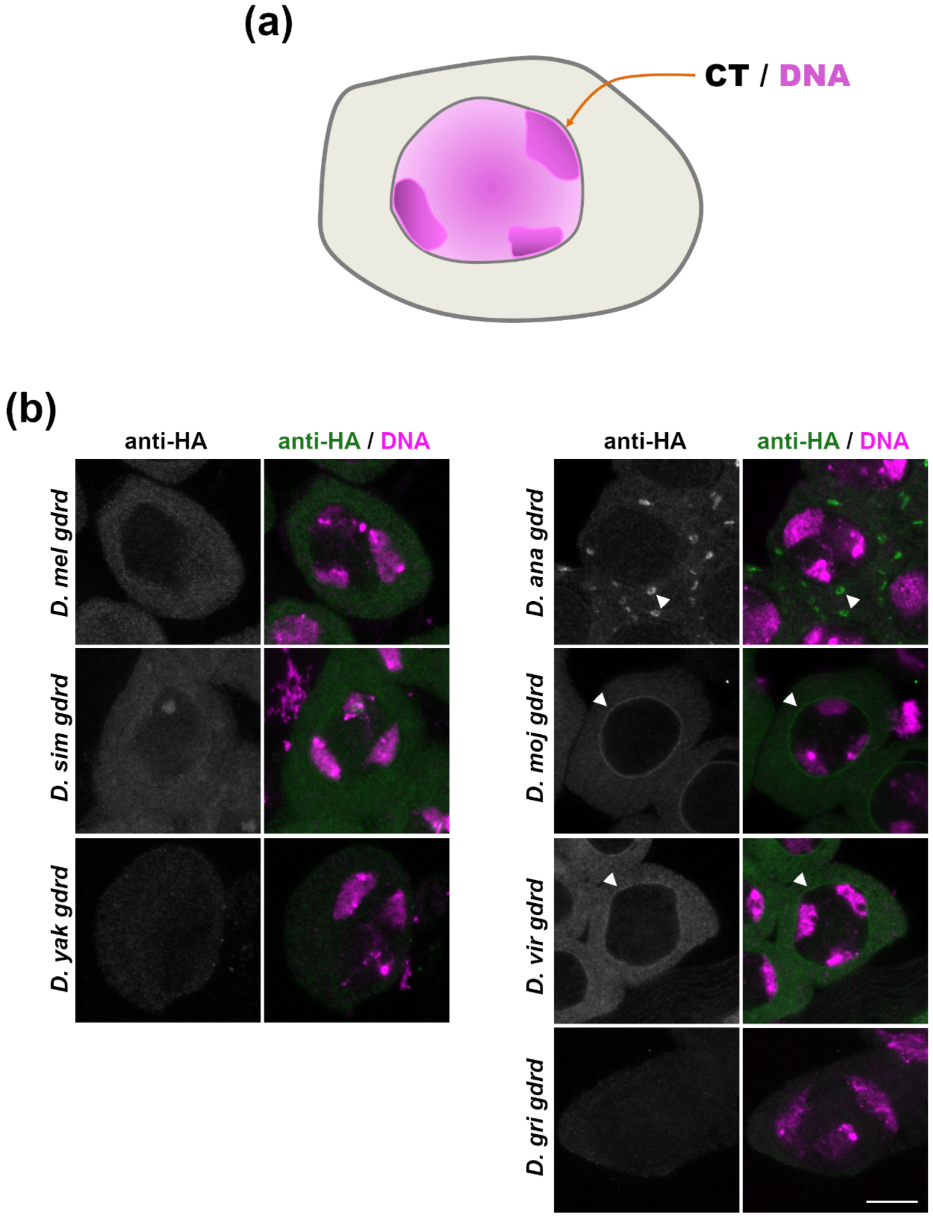


**Figure S12.** Gdrd orthologs exhibit divergent localization patterns during spermatocyte stage of spermatogenesis. a) Diagram: primary spermatocyte with characteristic chromosome territories (CT, magenta). b) Spermatocytes expressing Gdrd orthologs. *D. mel*, *D. sim*, and *D. yak* Gdrd proteins appear chiefly cytoplasmic, with substantial nuclear exclusion. The *D. ana* protein decorates round vesicular organelle structures, while the *D. moj* and *D. vir* proteins are enriched along the nuclear perimeter. Arrowheads mark examples of divergent localizations of orthologs in micrographs. Scale bar = 10 µm.

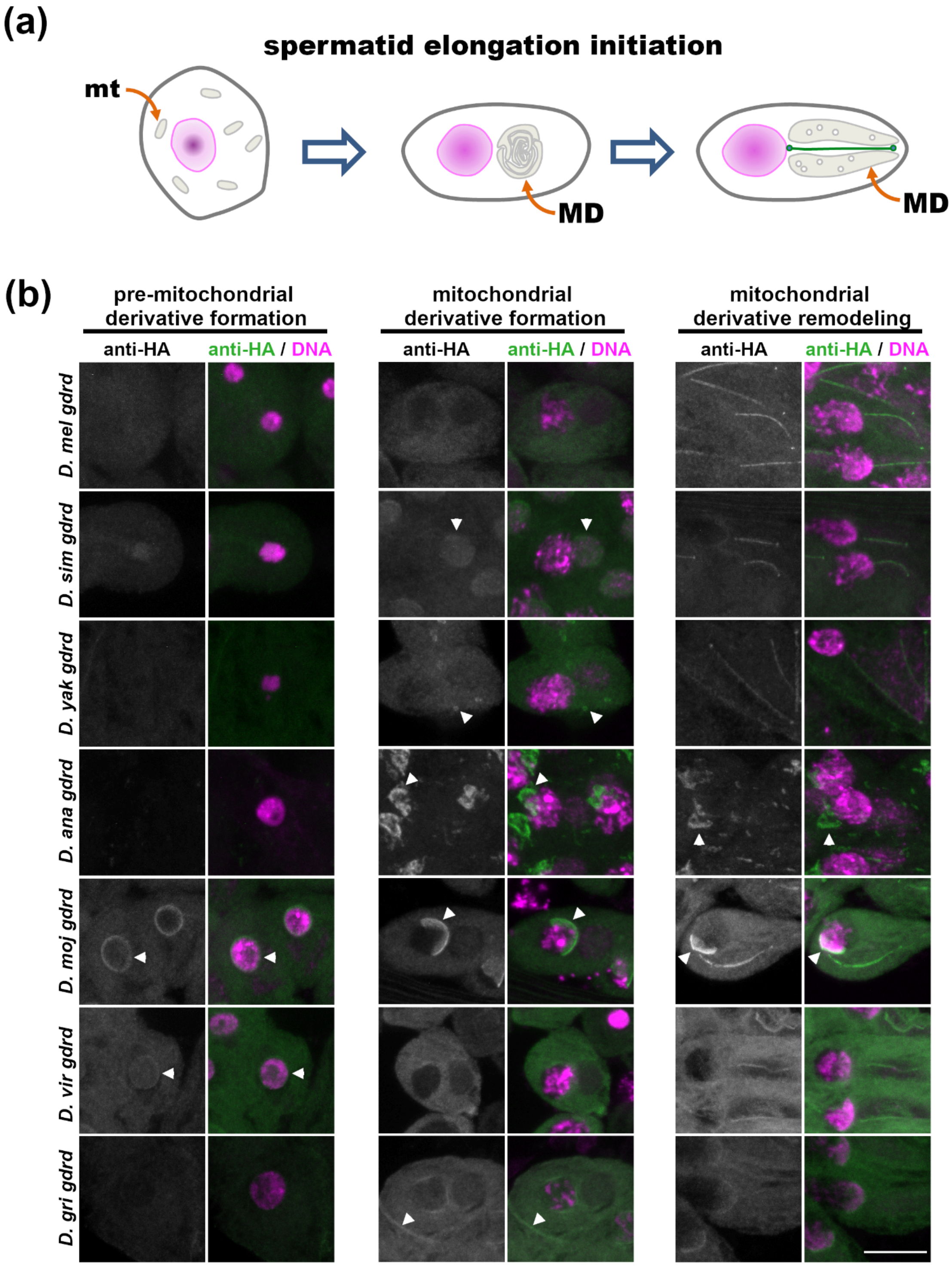


**Figure S13.** Gdrd orthologs exhibit divergent localization patterns during elongation initiation.

a) Diagram: mitochondrial dynamics drive spermatid elongation. In spermatids that have just completed meiosis, mitochondria (mt) are distributed throughout the cell. Spermatid elongation initiation begins with the fusion of cellular mitochondria into a round mitochondrial derivative (MD). Subsequent remodeling of the mitochondrial derivative drives spermatid elongation. b) Gdrd ortholog expression and subcellular localization was followed at three stages: (1) pre-mitochondrial derivative formation, (2) mitochondrial derivative formation, and (3) mitochondrial derivative remodeling. DAPI labels the nuclei (strong label) and the mitochondrial DNA (weak label). *D. mel* Gdrd is cytoplasmic until the onset of mitochondrial remodeling when it localizes to the basal body, axoneme, and transition zone. *D. sim* and *D. yak* Gdrd have a similar dynamic except that the proteins transiently localize to the mitochondrial derivative and round organellar structures (arrowheads), respectively, in spermatids undergoing mitochondrial derivative formation. The *D. ana* protein localizes to a structure reminiscent of the acroblast (arrowhead) in stages characterized by MD formation and remodeling. Notably, the *D. ana* protein shows weak localization to axonemes during mitochondrial derivative remodeling. At all three stages, *D. moj* and *D. vir* proteins have identical dynamics. The proteins decorate the nuclear perimeter in the pre-mitochondrial derivative stage. This uniform nuclear envelope distribution becomes restricted to the apical side at later stages of spermatid elongation initiation. In general, all *D. vir* ortholog localizations appear weaker when compared to the *D. moj* protein. The *D. gri* protein localizes to the plasma and nuclear membranes. Scale bar = 10 μm.

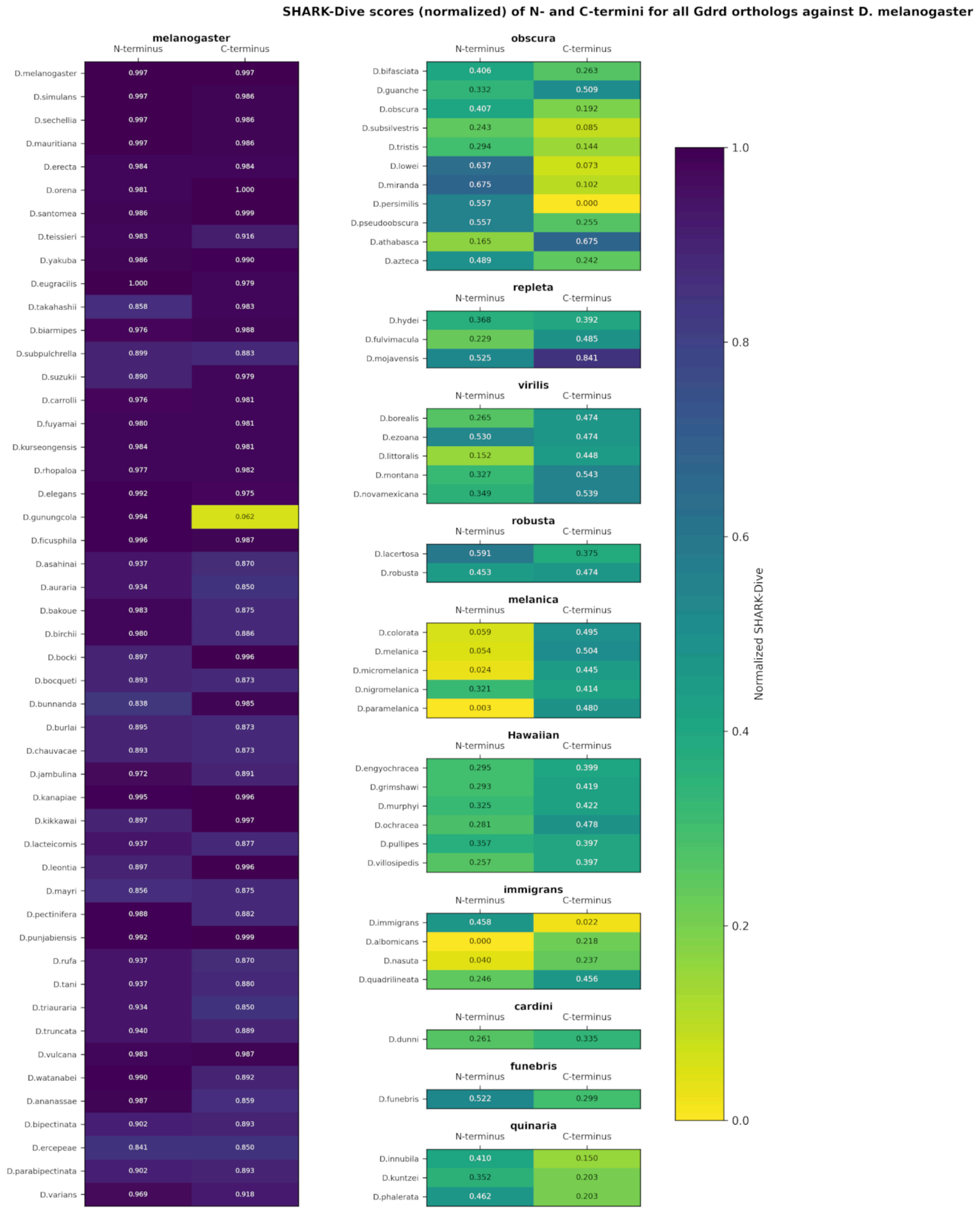


**Figure S14.** Normalized SHARK-Dive scores for the N- and C-termini of all identified Gdrd orthologs. The C-termini of *melanogaster* group species and the C-terminus of *D. mojavensis* likely share homology. Notably, the score for the *D. mojavensis* ortholog's

C-terminus is much higher than any other *Drosophila* subgenus species examined, indicating potential convergent evolution toward the *melanogaster* sequence within the *mojavensis* lineage.

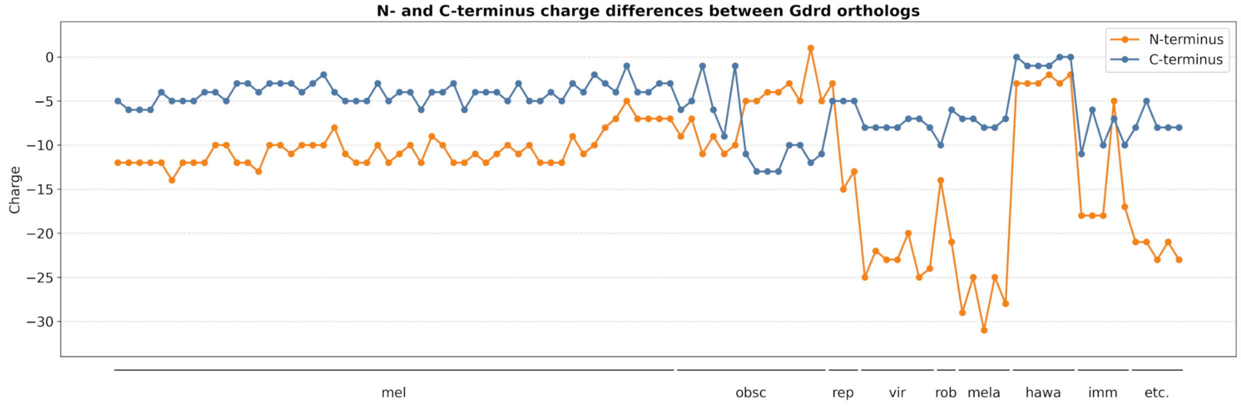


**Figure S15.** Overall ionic charges of orthologous Gdrd N- and C-termini. (mel, *melanogaster* species group; obsc, *obscura* species group; rep, *repleta* species group; vir, *virilis* species group; rob, *robusta* species group; mela, *melanica* species group; hawa, Hawaiian Drosophila; imm, *immigrans* species group; etc. includes species from the *cardini*, *funebris*, and *quinaria* species groups)

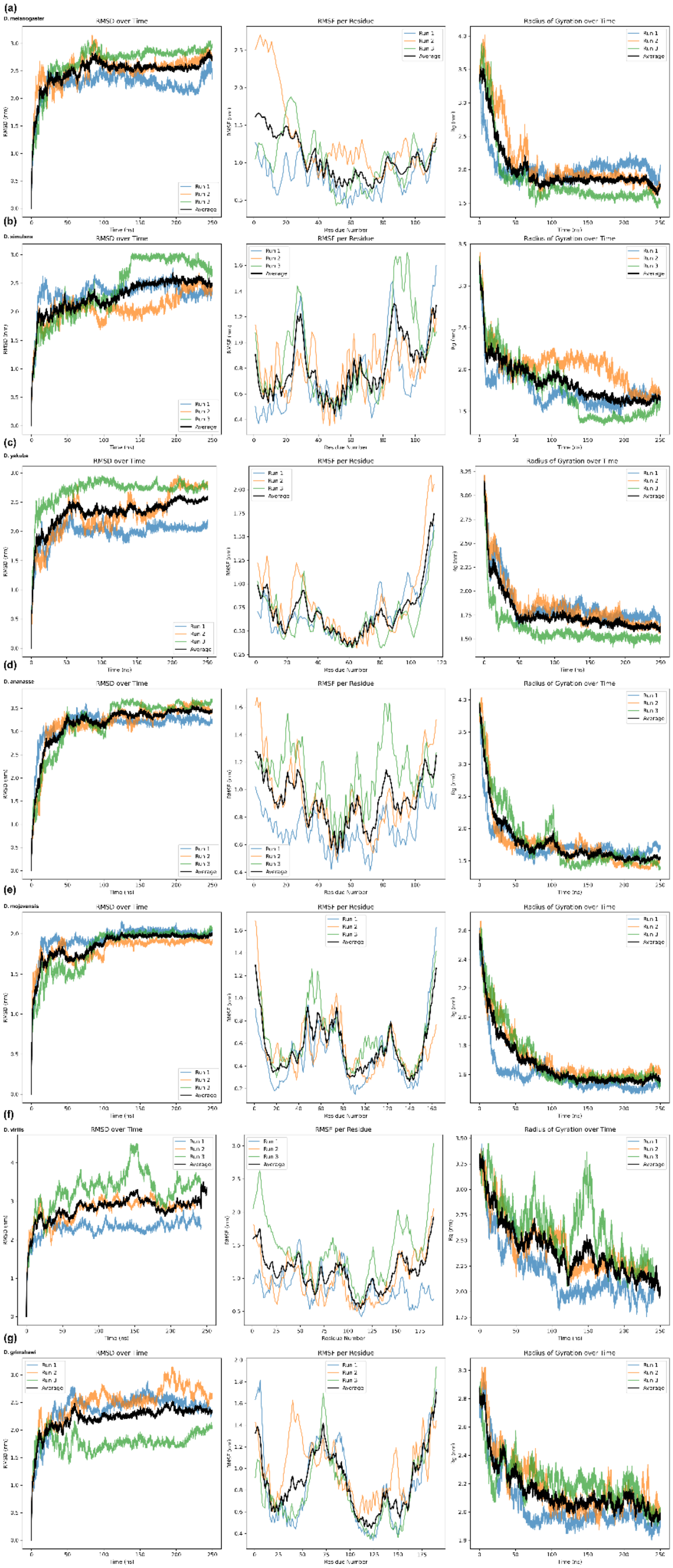
(a)
D. melanogaster
RMSD over Time
RMSF per Residue
Radius of Gyration over Time
(b)
D. simulans
RMSD over Time
RMSF per Residue
Radius of Gyration over Time
(c)
D. yakuba
RMSD over Time
RMSF per Residue
Radius of Gyration over Time
(d)
D. ananassae
RMSD over Time
RMSF per Residue
Radius of Gyration over Time
(e)
D. mojavensis
RMSD over Time
RMSF per Residue
Radius of Gyration over Time
(f)
D. virilis
RMSD over Time
RMSF per Residue
Radius of Gyration over Time
(g)
D. grimshawi
RMSD over Time
RMSF per Residue
Radius of Gyration over Time
Run 1
Run 2
Run 3
Average

**Figure S16.** Molecular dynamics (MD) simulations of Gdrd orthologs. Triplicate MD runs were performed for each Gdrd ortholog. Left column: backbone RMSD; middle column: $\alpha$-C RMSF; right column: radius of gyration (left to right). Each replicate is shown in color, and the average is shown in black. From top to bottom, the orthologs are: a) *D. melanogaster,* b) *D. simulans,* c) *D. yakuba,* d) *D. ananasse,* e) *D. mojavensis,* f) *D. virilis,* g) *D. grimshawi.*

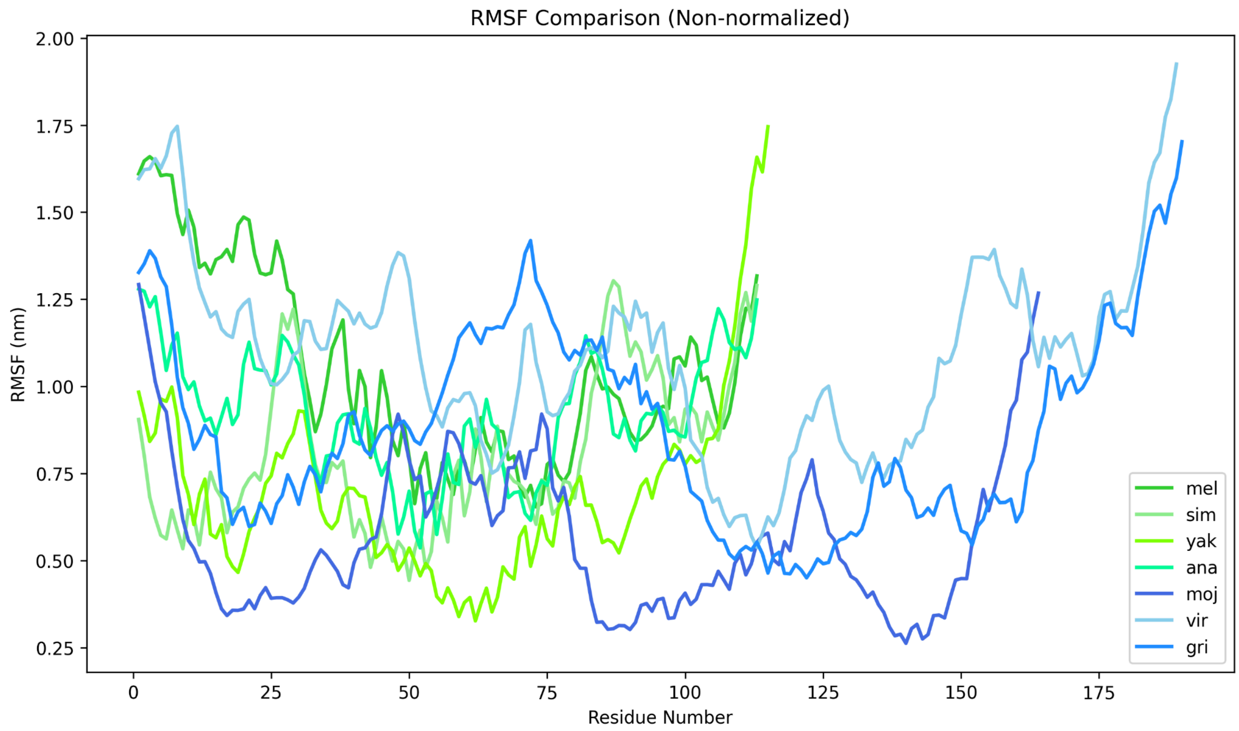


**Figure S17.** Per residue $\alpha$-C RMSF comparison for triplicate MD runs of Gdrd orthologs.
